# Irrational decisions reflect robustness constraints on value computations implemented by orbitofrontal circuits

**DOI:** 10.1101/2025.07.10.664081

**Authors:** Juliette Bénon, Mathias Pessiglione, Fabien Vinckier, Jean Daunizeau

## Abstract

Making good decisions is essential for survival and success, yet humans and animals often exhibit perplexing irrational decision-making whose biological origin remains poorly understood. Recent empirical and computational work suggests that altered computations in perceptual, motor and memory systems in the brain may arise from informational, metabolic or robustness constraints on their internal connectivity structure. However, whether and how such neurobiological constraints may have molded the architecture of decision systems (such as the orbitofrontal cortex) and eventually distorted decision-relevant computations, remains largely unknown. We first train cohorts of artificial neural nets to perform ten variants of rational decision-relevant computations. Those variants that operate under a specific option-encoding format exhibit most of the electrophysiological coding properties observed in orbitofrontal neurons of monkeys making decisions under risk. We then distort these neural nets’ internal wiring to reproduce monkeys’ irrational choices. This induces deterministic spillover interferences in decision-relevant computations that generalize across individuals, at both the behavioral and neural level. Importantly, although irrational nets do not seem to bring informational or metabolic benefits, they display enhanced tolerance to damage and noise when compared to their rational counterparts. This suggests that some forms of irrational behavior may be the incidental outcome of distal evolutionary pressure on the robustness of orbitofrontal circuits.

## Introduction

People and animals arguably act, in some circumstances, against their own interest. Why does irrational behavior persist, despite its potential costs to survival and fitness? Standard decision theory posits that rational decisions rely on estimating and comparing the expected value of each available alternative option in the choice set. Thus, irrational behavior may emerge from the covert mechanisms through which the brain constructs, represents, maintains or compares option values. Decades of work in human and non-human primates show that these computational processes involve a specific subset of brain systems, including – but not limited to – orbitofrontal (OFC), ventromedial (vmPFC) and dorsolateral prefrontal (dlPFC), as well as anterior cingulate (ACC), cortices^1–9^. While the relative contribution of these subsystems is not well understood, a robust finding across studies is that ventromedial and/or orbitofrontal neurons encode value, regardless of whether subjects are engaged in explicit decision-making or in the subjective evaluation of single options^10–13^. Accordingly, neuropsychological studies of brain-damaged patients demonstrate that lesions to the orbitofrontal cortex induce irrational value-based decisions without impairing other types of high-level cognitive processes^14–16^. This means that the effective rationality of decisions hinges on the integrity of OFC circuits. But even in the absence of anatomical lesions, value processing in the OFC is known to exhibit systematic distortions, which typically lead to overt irrational behavior. For example, value coding in the OFC is modulated by its pre-stimulus baseline activity^17,18^, adapts to the recent range of option values^19–21^, and depends on whether a given option is the status-quo alternative^22^ or is currently attended^23^. Given the behavioral biases that ensue, these results suggest that OFC circuits are organized in such a way that they process value-related information in a moderately, yet pervasively, suboptimal manner. In turn, this raises the basic question of why suboptimal value computations prevail in OFC circuits.

A common view is that seemingly irrational decisions may yield compensatory behavioral benefits. For example, inattention, overconfidence or optimism biases can be shown to be advantageous under the right circumstances^24–26^. However, an alternative assumption is that evolutionary pressure eventually selected for value computations that are “rational enough”, given the constraints that may act at the neurobiological level^27,28^. A prominent example is the energetic budget of neural circuits, which encompasses both physiological maintenance and activity-dependent firing costs^29,30^. The former costs typically restrains the total number of neurons in the brain^31,32^, which promotes neural coding strategies that minimize redundancy and/or maximize information transfer rates^33–36^. Interestingly, variants of such mechanisms explain value range adaptation effects in the OFC and the irrational behavior that ensues^20,28,37^. The latter costs induce metabolic budget constraints that are demonstrably tight. For example, the mitochondrial metabolic supply of neurons is actively restricted at the expense of circuit-level computational fidelity^38^, and a scarcity of external resources (e.g., food) eventually results in impaired neural processing^39^. This supports the idea that the brain has evolved so-called energy-efficient neural coding strategies that trade off computational precision for metabolic costs^40,41^. But theoretical work also emphasizes other types of tradeoffs that arise from demands on the robustness or fault-tolerance of neural circuits. A widely debated notion is that neural circuits must maintain their excitatory-inhibitory balance to ensure stability and/or homeostasis^42^. Another possibility, which is pervasive in biological systems, is the need to minimize vulnerability to localized damage^43,44^. Although direct empirical evidence for such a constraint on neural circuits is scarce, recent work indicates that frontal circuits that subtend, e.g. motor control and working memory, achieve tolerance to damage through architectural redundancy^45–47^. This is important because redundant systems are notoriously inefficient, from both an informational and energetic perspective^41,48^. In other words, OFC circuits may have evolved under multiple yet competing neurobiological constraints, whose impact on value-based decisions remains largely unexplored.

In principle, the fidelity of OFC value computations, its energy budget, and its biological robustness share a common determinant: its internal circuit wiring. From a neural net perspective, any developmental or evolutionary pressure of the sort discussed above will ultimately shape the architecture of OFC circuits in ways that distort value computations and compromise decision rationality^20,49^. This is, in fact, trivially observed in artificial neural net models of the OFC trained to perform candidate value computations while complying with these constraints (see Fig. 1). Critically however, the form of irrational behavior that emerges depends on both the nature of the constraint and the specific value computations the OFC is assumed to perform. This is because a given type of value computation requires a tailored neural net wiring, whose native compliance with the above constraints is largely arbitrary. Thus, the issues of identifying the nature of OFC value computations, the mechanisms through which they give rise to irrational behavior, and the biological constraints that may have molded the architecture of OFC circuits, are closely intertwined. In this work, we approach the problem from a computational perspective.

**Fig. 1.**
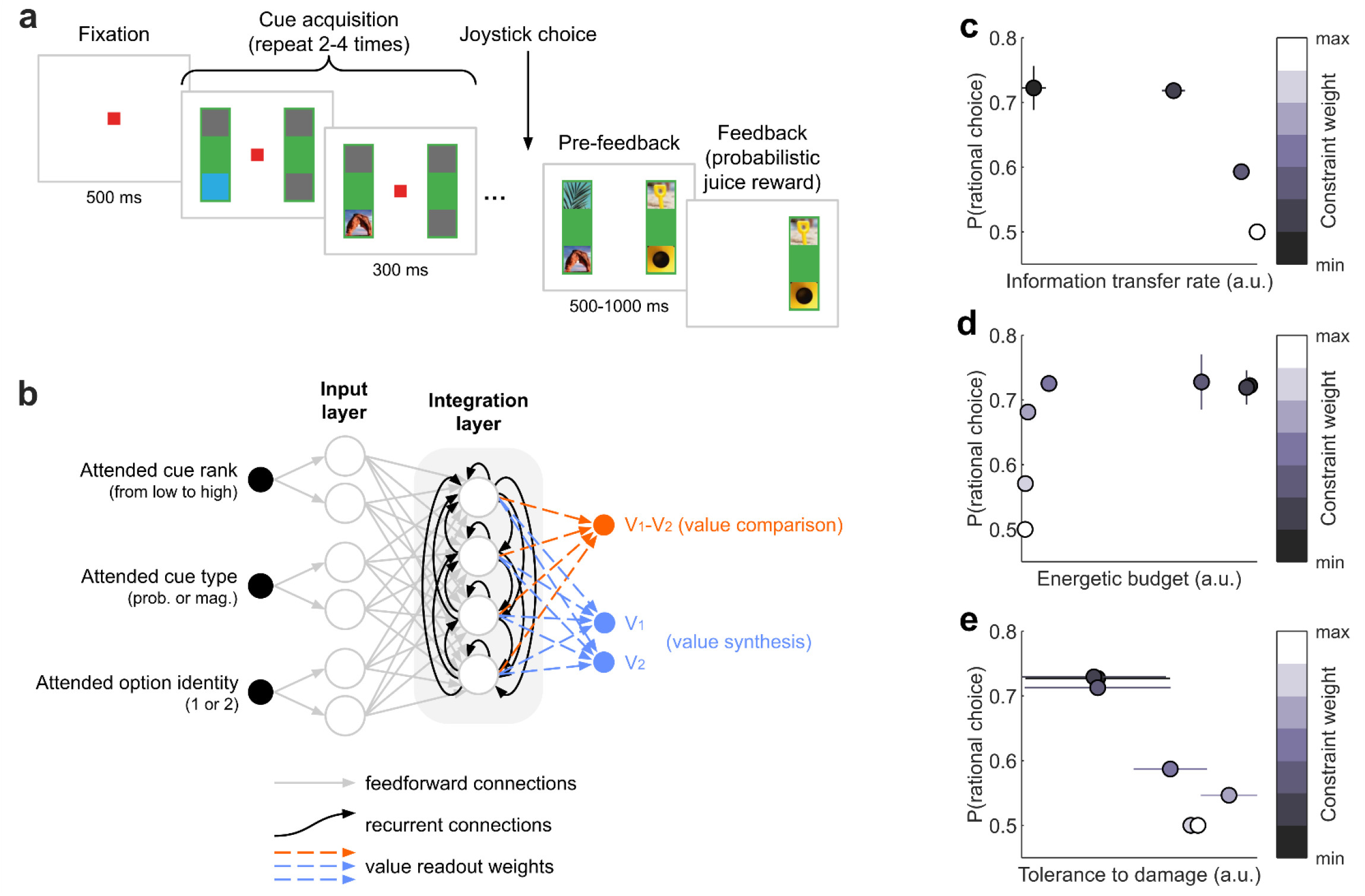
Decision task, neural net design, option values and impact of constraints. **a**, Task design. Adapted from Hunt et al., 2018^50^. At each trial, monkeys choose between two options presented on the left and right sides of a screen, each defined by the probability and prospective amount of a rewarding juice. Each decision cue (representing either the probability or the magnitude of the currently attended option) appears sequentially and then disappears. Monkeys can commit to a decision at any point after the second cue without necessarily sampling the remaining cues and are free to decide which cue to sample if they decide to continue the trial. **b**, Neural net architecture (see Methods). At each cue onset, the neural net’s inputs (back dots) encode the currently attended cue in terms of its three raw features (i.e., cue rank, cue type and option identity), while its outputs are the neural net’s current estimate of option values (value synthesis, blue dots) or value difference (value comparison, orange dot). Feedforward connections convey input features to a layer of input-specific units, which send their outputs to an input-integration layer. Importantly, the latter is equipped with recurrent connections that carry reentrant value computations elicited from previously attended cues. Note that both the identity of the attended option and value outputs can be encoded in distinct formats (see Methods). **c, d** and **e**, Neural net training under multiple constraints. Neural nets can be trained to optimize a compromise between rational value computations, on the one hand, and information transfer rate (**c**), energetic budget (**d**) or tolerance to damage (**e**), on the other hand (see Methods). Each dot depicts the average rate of rational choices (y-axis) and constraint adequacy (x-axis) for a given constraint compliance weight (color scale). Error bars indicate standard error. One can see that imposing stronger compliance with such constraints tends to compromise decision rationality, because the underlying network wiring eventually alters value computations.

We consider the paradigmatic case of binary decisions under risk, where the choice set consists of two alternatives, each defined by the probability and magnitude of prospective rewards. In principle, rational behavior amounts to choosing the option with the highest expected reward. Extensive electrophysiological recordings from OFC neurons are available in macaque monkeys performing this type of decision-making task. Here, we reanalyze an existing dataset in which decision cues (i.e. option-specific reward magnitude or probability) are revealed one at a time. This design provides a unique empirical estimate of the dynamics of information content in the OFC as value computations unfold over within-decision time^50^. In line with previous literature, we distinguish between two broad types of value computations: value *synthesis* and value *comparison*. The former implies that the OFC progressively integrates decision cues to compute the value of both options, which can be concurrently read out on possibly orthogonal activity subspaces of OFC neural ensembles^49,51,52^. The latter reduces to directly updating the value difference between the two options as a new decision cue becomes available^53,54^. Both value synthesis and value comparison can be implemented using one of five distinct neural encoding formats, which vary according to how cue inputs and value outputs are framed (see Results)^22,23^. Together, this yields a total of ten candidate scenarios regarding OFC value computations.

We first train recurrent neural nets (RNNs) to perform each candidate value computation in a rational manner, given arbitrary decision cue sequences. We then identify which, among the candidate types of value computations, yield accurate RNN models of the OFC. To do so, we compare the full set of recorded OFC neural responses with the activity patterns of simulated RNNs exposed to the same decision trials as those experienced by the monkeys. As we will see, this eventually selects two specific types of value computations, which effectively are idealized models of OFC networks that would have evolved without any metabolic or robustness constraint. We then distort the internal connectivity of these networks to reproduce monkeys’ irrational choices in the task (about 20% of all choices). As we will show, these distorted RNNs make behavioral and neural predictions that generalize across monkeys. Finally, we compare rational and irrational RNN models of the OFC, in terms of their compliance with energetic and robustness constraints. This enables us to identify which neurobiological constraint may have shaped OFC computations.

## Results

We took advantage of an open dataset of single unit activity recordings from the OFC, the dlPFC and the ACC of two macaque monkeys (n=189, 135 and 183 neurons respectively) engaged in value-based decision-making (22,618 trials in total)^50^. The task design is summarized on Fig. 1a (see also the Methods section). At each trial, monkeys chose between two options presented on the left and right sides of a screen, each defined by two cues (probability and prospective amount of a rewarding juice) that are revealed one at a time and could commit to a decision without necessarily sampling all cues. The subjective value profiles estimated from monkey choices (see Methods) indicate that they integrate both currently and previously attended cues, and are highly correlated with the rational value profile (Pearson correlation, Monkey F: *ρ* = 0.95; Monkey M: *ρ* = 0.93; all *p* < 10^-15^) with no significant correlation difference between monkeys (Steiger’s test, *p* = 0.19).

### Identifying value computations in the OFC

To begin with, we aimed to identify legitimate models of an idealized OFC network, which would have evolved without any metabolic or robustness constraint (and would thus perform rational value computations). To do this, we adopt a normative approach that obviates the need for empirical data in training models. In line with recent empirical work, we hypothesized that the OFC may implement one of two candidate decision-relevant computations: (i) computing the value of both options independently^51,55^ (“value synthesis”) or (ii) computing the difference between option values^9,56^ (“value comparison”). Each scenario can be implemented using recurrent artificial neural networks (RNNs), which operate under the exact same conditions as monkeys in the task. In particular, RNNs access cues sequentially and in an encoding format that specifies attribute type and rank as well as option identity (see below). At each cue onset, these inputs are sent to a first hidden layer (cue-encoding), whose units feed their output forward to a second hidden layer (cue-integration), from which the RNN’s value outputs are readout (see Fig. 1b and Methods). The integration layer relies on internal recurrent connections to combine currently and previously sampled cues, and progressively update its ongoing within-trial computations^49^. Importantly, both value synthesis and comparison also require specifying how options are identified, which is debated in the existing literature. The OFC may do so based on, e.g., spatial location^57^ (left vs. right), temporal order^49^ (first vs second), or attentional focus^23^ (attended vs. unattended). In principle, both OFC inputs (decision cues) and outputs (option values or value difference) may encode option identity in a different format, irrespective of whether the OFC operates value synthesis or comparison. This resulted in ten cohorts of RNNs (two types of value computations combined with five input-output format variations), which we train on rational option values (see Methods). This is not trivial, since it requires RNNs to maintain a memory trace of previously attended cues, while remaining invariant to the order in which cues are presented. Importantly, each cohort gathers 1,000 RNN instances that sample the manifold of admissible wirings following random weight initializations.

Once trained, these RNN models can be used to simulate neural population dynamics that implement rational value computations during decision trials. If triggered with the sequences of cues that monkeys attend in the decision task, this provides value readouts and activity patterns that can be compared to observed choices and OFC recordings (see Fig. 2). When probed at monkeys’ decision onset time, these rational RNNs correctly predict 79% ± 3×10^-3^ (mean ± standard error) of choices (monkey F: 77% ± 4×10^-3^, monkey M: 80% ± 4×10^-3^). Crucially, although all rational RNNs yield identical decisions, their internal representations are different (see Supplementary Materials). We thus asked whether any of these RNN cohorts capture key aspects of OFC neural representation geometry, despite not having been exposed to neural recordings during training. To test this, we replicated the two types of analysis conducted by Hunt and colleagues^50^ on single units’ recordings, which we also performed on the RNNs’ integration layer (see Fig. 2a). We first ran a representational similarity analysis at first cue onset, building representational dissimilarity matrices (RDMs) by correlating population activity vectors in response to all possible cues (see Methods). In brief, RDMs identify which cue features elicit discriminable response patterns across neurons when only a single cue is available. To track neural representation geometry at all stages of decision trials, we also quantified whether and how inter-neuron differences in their sensitivity to current and past cue ranks are preserved across cue onset times (cf. cross-correlation matrices or CCMs; see Methods). One can think of RDMs and CCMs as two distinct summary statistics of the representational geometry of distributed neural systems, with complementary properties (see Fig. 2b). We then derived the two ensuing neural distance metrics by comparing OFC neurons and RNN units at each stage of the training process (see Fig. 2c). Note that even untrained – i.e. random – RNNs exhibit some degree of neural similarity with the OFC, because they respond to value-relevant input cues. Untrained RNNs thus effectively provide the distribution of neural distances under the null hypothesis. Now, when being trained to perform a specific value computation, RNNs modify their representational geometry and hence their neural distance to the OFC. In line with previous work on visual and language systems in the brain^58,59^, we considered that legitimate idealized RNN models of the OFC are those RNN cohorts that significantly decrease both neural distance metrics as a result of training.

**Fig. 2.**
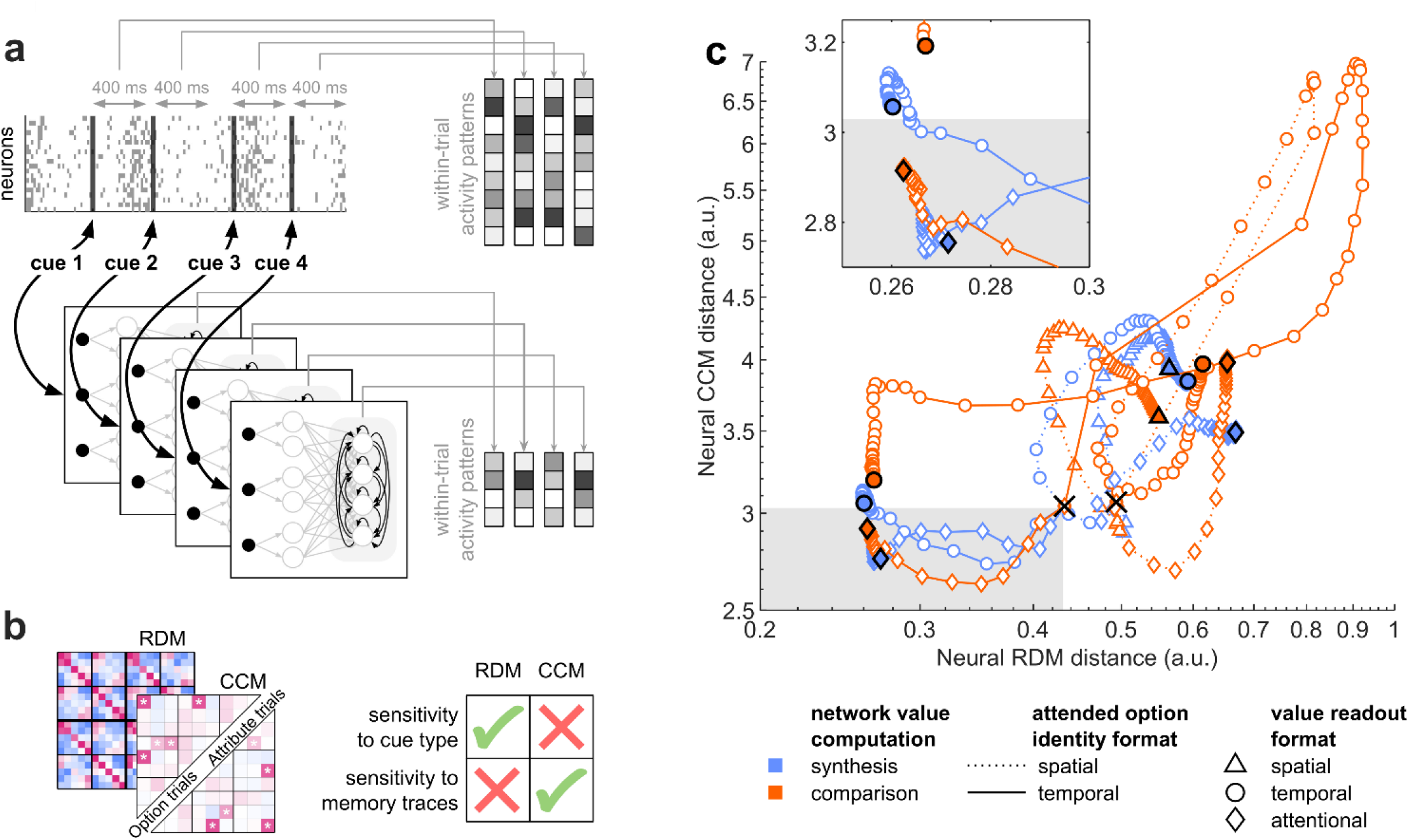
Selection of candidate idealized RNN models of the OFC. **a**, Schematic procedure for extracting within-trial activity patterns from OFC neural recordings and RNN models. At each trial, a sequence of cues is sampled until a choice is triggered. Within-trial activity patterns of OFC neurons are constructed as the average firing rate of each neuron, from 100 ms to 500 ms after each cue onset. Similarly, within-trial activity patterns of RNN models are the activation strengths of integration units in response to each cue. **b**, Summary of neural distance metrics for comparing activity patterns of OFC neurons and RNN models. RDMs quantify how dissimilar evoked activity patterns are for any pair of cues (2×2×5 = 20 possibilities at first cue onset). CCMs quantify the similarity of profiles of neural sensitivity to present and past cue ranks. Although CCM-based distance metrics are insensitive to cue type (probability or magnitude), they quantify potential internal memory traces about previously sampled cues. Full RDM and CCM summary statistics for all monkeys and brain regions can be eyeballed in the Supplementary Materials. **c**, Average neural distance trajectories between OFC and RNN cohorts (over 1,000 RNN instances), computed using either RDMs (x-axis) or CCMs (y-axis) metrics. Black crosses indicate the initial (random) state of RNN cohorts, black triangles/dots/diamonds denote their final rational state. Intermediary points show the neural distance at various stages of RNN training (from 0% to 100%, by steps of 2%), where color, line style and marker type indicate the type of computation (value synthesis vs value comparison), the identity format of the attended option (spatial vs temporal), and the value readout format (spatial vs temporal vs attentional), respectively. The gray shaded area indicates the two-dimensional region that falls below the 95% confidence interval of both pre-training neural distances.

Strikingly, all RNN cohorts that rely on the spatial (left versus right) encoding of the attended option’s identity tend to increase both neural distances to OFC neurons as training unfolds. Among the remaining four cohorts, those that compute option values using a temporal format (first versus second) eventually shorten their RDM distance, but at the cost of compromising their CCM distance. Ultimately, only those two RNN cohorts that encode the attended option identity using the temporal format (first versus second) while computing option values in the attentional format (attended versus unattended) significantly shorten both neural distances during training. These differ in terms of the type of value computation: one RNN cohort performs value synthesis (CCM distance, paired two-sided t-test: *t*(999) = 9.5, *p* < 10^-15^, Cohen’s *d* = 0.30; RDM distance: *t*(999) = 44, *p* < 10^-15^, Cohen’s *d* = 1.39), whereas the other performs value comparison (CCM: *t*(999) = 4.6, *p* = 5×10^-6^, Cohen’s *d* = 0.14; RDM: *t*(999) = 48, *p* < 10^-15^, Cohen’s *d* = 1.52). Although we cannot yet arbitrate between these two scenarios, we have clearly narrowed the set of plausible models of the idealized OFC network. In the remainder of the paper, we focus on these two RNN cohorts (some extended results for all model variants are shown in the Supplementary Materials).

At this point, we asked whether and how models of the idealized OFC network can be modified to explain monkeys’ irrational behavior. We thus retrained the selected rational RNNs to predict monkeys’ choices in the task, of which about 20% are irrational (Monkey F: 21% ± 3×10^-2^; Monkey M: 19% ± 3×10^-2^). To preserve the interpretability of RNN value computations while allowing for, e.g., nonlinear interferences and spill-over effects between attended cues, RNNs were initialized with their trained rational weights, and retraining was restricted to recurrent connections within the integration layer. When probed at monkeys’ decision onset time, retrained RNNs achieve an average choice-prediction accuracy of about 83% ± 1×10^-2^ (monkey F: 80% ± 5×10^-2^, monkey M: 84% ± 2×10^-2^), significantly outperforming rational models (synthesis models, paired two-sided t-test: *t*(1999) = 154, *p* < 10^-15^, Cohen’s *d* = 3.44; comparison models: *t*(1999) = 174, *p* < 10^-15^, Cohen’s *d* = 3.88; see Fig. 3e). Moreover, models trained on one monkey make choice predictions on the other monkey that still significantly outperform rational RNNs (synthesis models, paired two-sided t-test: *t*(1999) = 25, *p* < 10^-15^, Cohen’s *d* = 0.55; comparison models: *t*(1999) = 22, *p* < 10^-15^, Cohen’s *d* = 0.49; see Fig. 3e). This means that retrained RNNs capture hidden deterministic mechanisms underlying irrational behavior that generalize across trials and individuals.

**Fig. 3.**
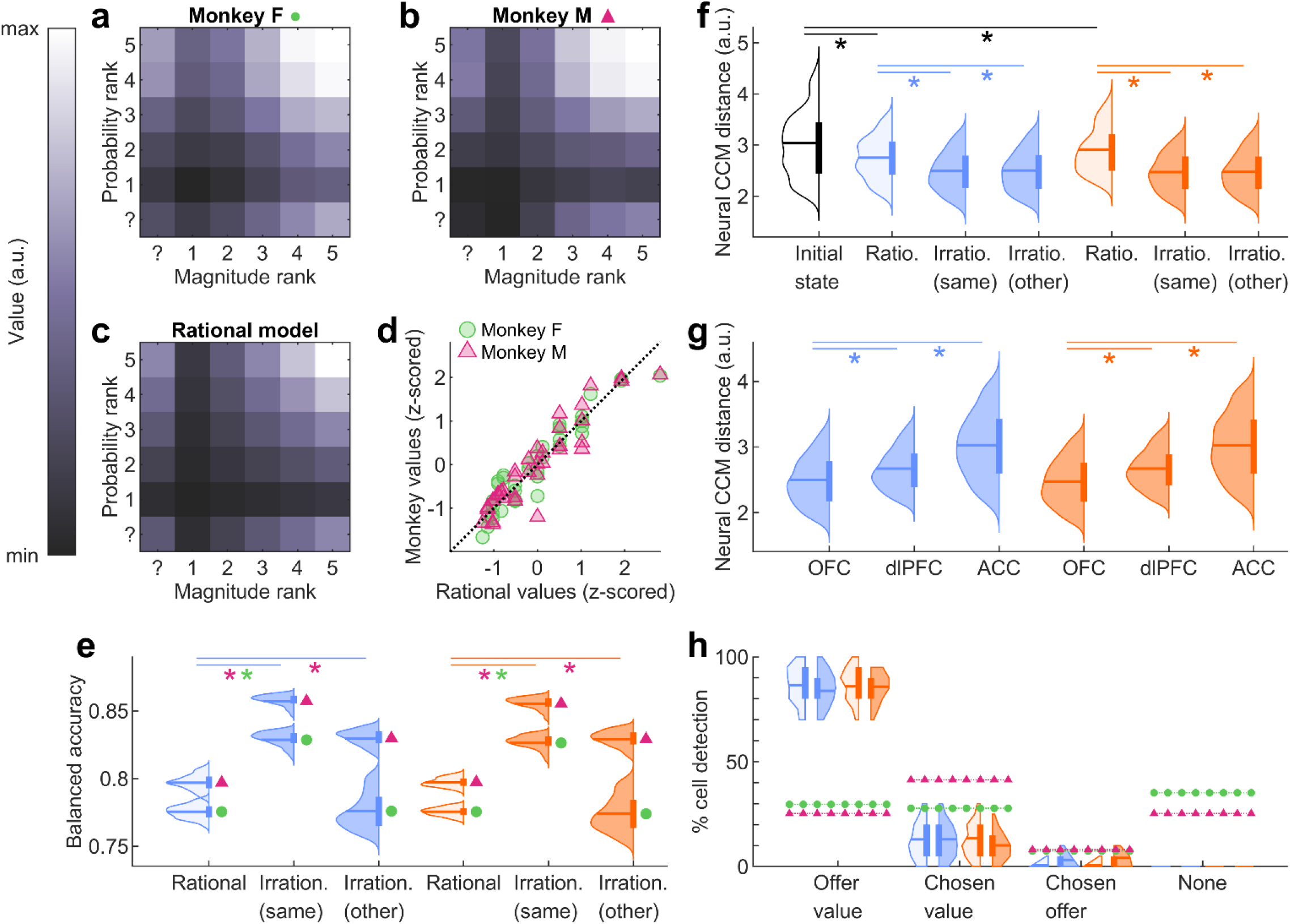
Behavioral and neural realism of candidate RNN models of the OFC. **a** and **b**, Estimated option values (see Methods) for monkey F (**a**) and M (**b**) are shown as a function of reward’s magnitude rank (x-axis) and probability rank (y-axis). **c**, Rational options values are defined as the expected reward, i.e. the product of reward magnitude and probability. Note that when an attribute is unknown (cf. question mark), the rational model replaces it with its a priori expected rank. **d**, Monkeys’ estimated values (y-axis) are plotted against rational values (x-axis). Each point represents a specific combination of probability and magnitude ranks, including cases where one or both attributes are unknown. **e**, Balanced accuracy for predicting monkey choices (green circle: monkey F, pink triangle: monkey M), under each candidate model (blue: value synthesis, orange: value comparison). Lighter distributions correspond to rational models, darker distributions to irrational models, and right-most distributions represent irrational models trained on one monkey and tested on the other. Within each violin plot, the horizontal line denotes the mean, and the thicker vertical line represents the interquartile range (25^th^ – 75^th^ percentile). Asterisks indicate significant differences, with p-value < 0.005. **f**, Neural CCM distance between models and the OFC. The white distribution corresponds to random RNN initializations (identical for both RNN cohorts). **g**, Neural CCM distance (averaged across monkey-specific distances) between irrational models and the OFC, the dlPFC and the ACC. **h**, Percentage of units classified as *offer value, chosen value* or *chosen option* cells, in rational and irrational RNN models and in recorded OFC neurons (green circle: monkey F, pink triangle: monkey M), at the time of choice.

Although we have leveraged the flexibility of RNNs to capture irrational choices, we have not yet demonstrated that irrational RNNs are accurate models of actual OFC computations. Remarkably, their neural CCM distance to the OFC decreases even further compared to their rational counterparts (synthesis models, paired two-sided t-test: *t*(1999) = 17, *p* < 10^-15^, Cohen’s *d* = 0.38; comparison models: *t*(1999) = 29, *p* < 10^-15^, Cohen’s *d* = 0.64). Furthermore, this improvement generalizes across monkeys, as shown when evaluating the neural distance of retrained RNNs to the other monkey (synthesis models, paired two-sided t-test: *t*(1999) = 16, *p* < 10^-15^, Cohen’s *d* = 0.36; comparison models: *t*(1999) = 27, *p* < 10^-15^, Cohen’s *d* = 0.61; see Fig. 3f). This is despite yielding activity patterns that accurately predict inter-individual differences in CCM matrices (see Supplementary Materials). However, one may argue that informing RNN models about monkeys’ actual choices may have facilitated the resemblance to any brain system that contributes to behavioral control in the task, thus challenging the anatomical specificity of our results. To address this point, we also computed the neural distance of retrained RNNs to dlPFC and ACC neurons. We first checked that empirical summary statistics of neural information geometry vary more across brain regions than across monkeys (see Supplementary Materials). We then compared neural distances across brain regions; we found that retrained RNNs were significantly closer to the OFC than to the dlPFC and the ACC (synthesis and comparison models, paired two-sided t-test: *t*(1999) > 29, *p* < 10^-15^ and Cohen’s *d* > 0.65 for all comparisons between areas, see Fig. 3g).

One may also ask whether selected RNNs exhibit the established mixed selectivity of OFC neurons. In line with the existing literature^57,60^, we attempted to classify units according to three distinct response profiles (see Methods): *option value cells*, which encode the value of a single option; *chosen option cells*, which encode the binary identity of the chosen option; and *chosen value cells*, which encode the value of the chosen option. As expected, we found that the trial-by-trial variations of OFC neurons’ firing rate at the time of choice can be matched to one of the three response profiles. Importantly, this is also the case for integration units of selected RNNs, albeit with a slight over-representation of *offer value* units (see Fig. 3h). Note that this gap is much reduced if we remove the OFC neurons that do not match any of those response profiles (about 30% of OFC neurons).

Together, these findings suggest that the selected RNNs perform value computations that are realistic, from both a behavioral and neural standpoint. In particular, adjusting the RNNs’ internal wiring to account for irrational choices yields both accurate and specific predictions of the OFC’s representational geometry. This completes the identification of idealized (rational) and actual (irrational) neural net models of value computations in the OFC.

### Characterizing irrational interferences in value computations

We still lack a computational explanation for why monkeys exhibit irrational behavior in the task. Thus, we now seek to characterize the systematic distortions in cue processing that give rise to irrational choices. First, we quantified potential nonlinear interference effects across decision cues. Recall that, by assumption, rational choices should be solely driven by the informational content of decision cues and thus remain invariant with regard to cue presentation order. In contrast, irrational interference effects would manifest as variability in RNNs’ value outputs across random permutations of cue presentation order, all else being equal. We thus performed systematic simulations of selected RNNs, quantifying the standard deviation of value outputs across permuted cue presentation orders, for all possible combinations of two, three or four cues (see Methods).

By construction, rational RNN models exhibit almost no variability. Thus, irrational RNNs exhibit significantly stronger interference effects than their rational counterparts (paired two-sided t-test at each time step: all *t*(1999) > 64, *p* < 10^-15^ and Cohen’s *d* > 1.44). Importantly, interference effects increase as within-trial decision time unfolds (paired two-sided t-test within each cohort between step 2 and step 4: both *t*(1999) > 227, *p* < 10^-15^, Cohen’s *d* > 5.08; see Fig. 4b and Fig. 4c). This suggests that systematic perturbations in sequential cue processing accumulate over time. Accordingly, monkeys’ choices become more irrational – i.e. less consistent with their average preferences (cf. Fig. 3a and Fig. 3b) – as decision time unfolds (Monkey F, paired two-sided t-test between pairs of steps: all *t*(24) > 3.3, *p* < 3×10^-3^, Cohen’s *d* > 0.68; Monkey M, step 3 vs. step 4: *t*(29) = 13, *p* = 2×10^-13^, Cohen’s *d* = 2.45; see Fig. 4d). One may argue that this accumulating interference effect may only be apparent, because decisions that monkeys trigger later in time tend to be more difficult (see Supplementary Materials). To control for the effect of decision difficulty, we regressed irrational choice rates onto the absolute subjective value difference, across trials. Reassuringly, the residuals of this regression still increase as decision time unfolds (Monkey F, two-sample two-sided t-test between pairs of steps: all *t* > 3.4, *p* < 7×10^-4^, Cohen’s *d* > 0.07; Monkey M, step 3 vs. step 4: *t*(12830) = 5.2, *p* = 2×10^-7^, Cohen’s *d* = 0.09; see Fig. 4e). This means that monkeys’ rationality deteriorates with decision time beyond what can be expected from decision difficulty alone.

**Fig. 4.**
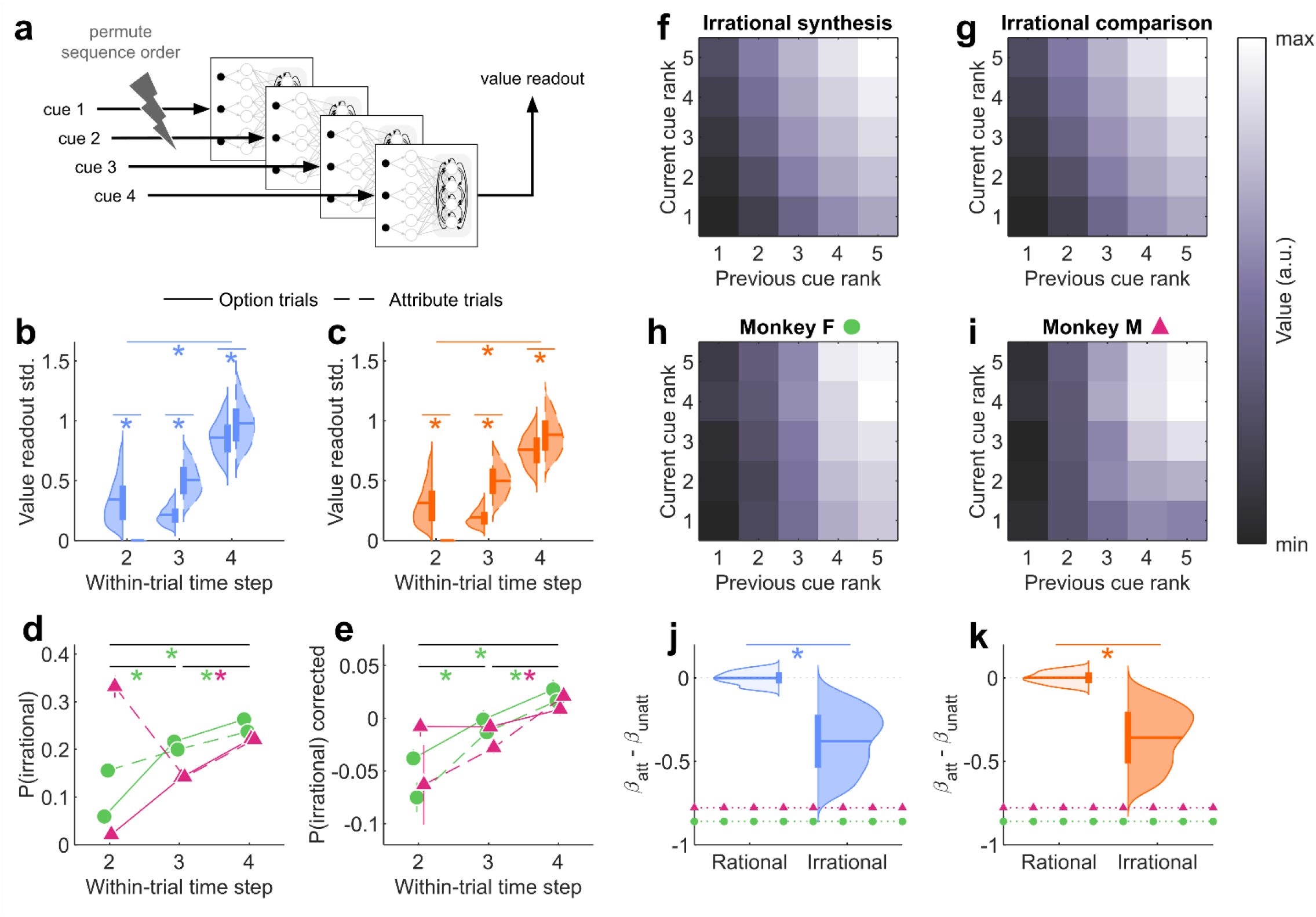
Interference mechanisms in irrational models and monkeys. **a**, Schematic procedure for evaluating the RNNs’ sensitivity to cue presentation order. **b**, Standard deviation of the irrational value synthesis RNNs’ outputs in response to random permutations of cue sequence orders (y-axis), as a function of cue onset times (x-axis) during option trials only (solid line) or attribute trials only (dashed line). Asterisks indicate p-value < 0.005. **c**, Same format as panel **b**, but for irrational value comparison RNNs. **d**, Rate of monkeys’ irrational choices (y-axis; green circle: monkey F, pink triangle: monkey M), as a function of cue onset time, for both option (solid) and attribute (dashed) trials. Vertical error bars show the standard error. Asterisks indicate that the difference between time steps (averaged over both trial types) are significant, with p-value < 0.02. **e**, Average residual irrational choice rate, once decision difficulty has been regressed away (same format as panel **d**). **f**, Average value output (color scale) of irrational value synthesis RNNs, as a function of the rank of both previously (x-axis) and currently (y-axis) attended cues (see Methods). **g**, Same format as panel **f**, but for irrational value comparison RNNs. **h** and **i**, Same format as panel **f**, but for both monkeys (h: monkey F, i: monkey M). **j**, Difference between the contributions of cue ranks to the attended pseudo-value (attended cue minus unattended cue; see Methods), for both rational (light) and irrational (dark) variants of value synthesis RNNs. The asterisk denotes a significant difference between rational and irrational RNNs, with p-value < 0.01. Colored dotted lines indicate the attended/unattended cue contribution difference for both monkeys. **k**, Same format as panel **j**, but for value comparison RNNs.

A possibility is that cue traces within the RNNs’ integration layer may leak into one another, either across options or across attributes. To investigate this, we separated *option trials* – where the second cue reveals the missing attribute of the same option as the first cue – from *attribute trials* – where the second cue reveals the same attribute as the first cue, but for the other option. At the second cue onset, interference effects are significantly stronger in option trials than in attribute trials, for both RNN types (paired two-sided t-test within each cohort: both *t*(1999) > 64, *p* < 10^-15^, Cohen’s *d* > 1.45). This is also the case for monkey F, based on residual irrational choice rates (Monkey F, two-sample two-sided t-test: *t*(1169) = 2.4, *p* = 2×10^-2^, Cohen’s *d* = 0.14; Monkey M: *t*(321) = 1.6, *p* = 0.11, Cohen’s *d* = 0.18; see Fig. 4e). This suggests that cue interference effects are more pronounced within options – i.e. across attributes – than across options. Thus, when attending a given option, we expect the integration of previously and currently attended cues to be asymmetrical, above and beyond differences induced by the type of information that they convey – i.e. reward probability vs magnitude. To test this, we quantified a pseudo-value profile as a function of both previously and currently attended cue ranks, irrespective of cue types (see Methods). In contrast to rational RNNs, irrational RNNs output pseudo-values that are mostly influenced by the previously attended cue (see Fig. 4f and Fig. 4g). When quantified in terms of the relative contribution of the unattended versus attended cue ranks (see Methods), we find that the asymmetry is significantly stronger in retrained RNNs than in rational RNNs (paired two-sided t-test within each cohort: both *t*(1999) > 220, *p* < 10^-15^,Cohen’s *d* > 4.96; see Fig. 4j and Fig. 4k). This asymmetry is also significantly present in monkeys’ choices (Monkey F, one-sample two-sided t-test: *t*(23) = -18, *p* < 10^-14^, Cohen’s *d* = 3.72; Monkey M: *t*(28) = -17, *p* < 10^-15^, Cohen’s *d* = 3.24; see Fig. 4j and 4k). Together, these results suggest that previously attended cues leave a persisting value trace that partly resists novel value-relevant information, yielding accumulating perturbations in value computations.

### Comparing biological constraint compliance in rational and irrational RNNs

In contrast to irrational RNNs, rational RNNs are idealized neural net models of OFC circuits that would have evolved in the absence of energetic or robustness constraints. We now ask whether the wiring peculiarities of irrational RNNs may bring some form of biological advantage that may have overcompensated the behavioral suboptimality that they induce.

First, we compared rational and irrational RNNs in terms of the metabolic cost of sustaining their respective structures. Since action potentials are a major source of energetic consumption in the brain^61^, we quantified metabolic cost in terms of the average network activity (see Methods). However, we found no significant difference in energetic budget between rational and irrational RNNs (synthesis models, paired two-sided t-test: *t*(1999) = 2.0, *p* = 0.04, Cohen’s *d* = 0.05; comparison models: *t*(1999) = 1.2, *p* = 0.25, Cohen’s *d* = 0.03; see Fig. 5a).

**Fig. 5.**
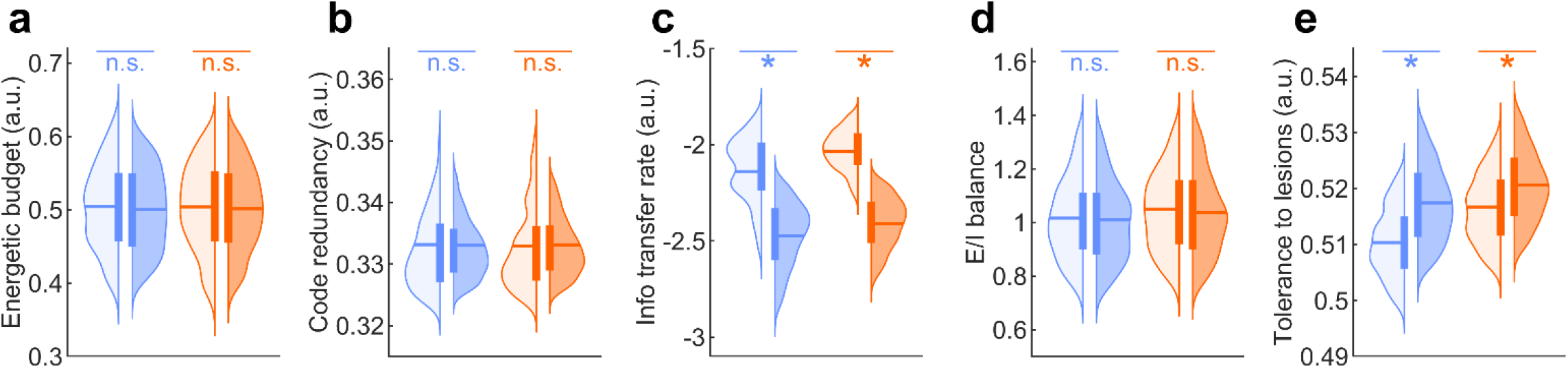
Potential biological benefits of irrational circuits. For all panels, asterisks indicate a significant difference between rational (light) and irrational (dark) RNNs (blue: value synthesis, orange: value comparison), with p-value < 0.005. **a**, Metabolic cost, measured as the average network activity, over all trials, trial steps, and units. **b**, Neural code redundancy, measured as the average co-activation probability over all integration units pairs. **c**, Information transfer rate, measured as the entropy of units response outputs. **d**, Excitatory-inhibitory balance, measured as the relative proportion of positive and negative connection weights. **e**, Tolerance to damage, measured as the retained rate of rational choice from 10% to 50% of lesioned units.

Second, we took inspiration from other variants of efficient coding models, which rather suggests that brain circuits with limited neural resources self-organize to either minimize code redundancy or maximize information transfer rate. We quantify these in terms of the average rate of units’ co-activation across all possible units pairs^62,63^ and the entropy of units’ response outputs^20,64^, respectively (see Methods). We found no significant difference in code redundancy (synthesis models, paired two-sided t-test: *t*(1999) = 0.4, *p* = 0.68, Cohen’s *d* = 0.01; comparison models: *t*(1999) = -0.8, *p* = 0.42, Cohen’s *d* = 0.02; see Fig. 5b). Interestingly however, we found that irrational RNNs exhibit significantly lower information transfer rate than their rational counterparts (synthesis models, paired two-sided t-test: *t*(1999) = 51, *p* < 10^-15^, Cohen’s *d* = 1.13; comparison models: *t*(1999) = 82, *p* < 10^-15^, Cohen’s *d* = 1.82; see Fig. 5c). This implies that the wiring peculiarities that yield irrational behavior also incur an additional cost in terms of information transfer rate.

Third, we reasoned that irrational circuits may benefit from better robustness. We start with excitatory-inhibitory balance, which would ensure stability and/or homeostasis^42^. However, we found no significant difference in the relative proportion of negative and positive connection weights (see Methods) between rational and irrational RNNs (synthesis models, paired two-sided t-test: *t*(1999) = 1.2, *p* = 0.23, Cohen’s *d* = 0.03; comparison models: *t*(1999) = 2.3, *p* = 0.02, Cohen’s *d* = 0.05; see Fig. 5d).

We then reasoned that irrational circuits may simply be more tolerant to external perturbations such as damage or noise. To test this, we simulated random virtual lesions of RNN integration units and measured the retained rate of rational choices. As expected, rational choice rate monotonically decreases when the fraction of lesioned units increases, for all types of models. Thus, we quantify tolerance to damage in terms of the rational choice rate averaged within a range of lesion sizes (from 10% to 50% of integration units; see Methods). We find that irrational RNNs exhibit significantly stronger tolerance to damage than their rational counterparts, irrespective of value computations (synthesis models, paired two-sided t-test: *t*(1999) = -27, *p* < 10^-15^, Cohen’s *d* = 0.61; comparison models: *t*(1999) = -16, *p* < 10^-15^, Cohen’s *d* = 0.36; see Fig. 5e). Interestingly, perturbing OFC computations with neural noise or imposing virtual disconnections within the integration layer yield qualitatively identical results (see Supplementary Materials).

## Discussion

In this work, we asked whether irrational behavior may be explained by distal constraints that act on the neurobiology of brain decision-making systems. First, we adopted a normative approach to identify models of an idealized OFC network, which would have evolved without any metabolic or robustness constraint. We found that only a specific subset of candidate RNNs reproduces the representational geometry of the OFC – specifically, those that receive inputs encoding option identity in a temporal format (first vs. second option), while computing option values or value difference in an attentional format (attended vs. unattended option). We discuss this finding below. Second, we retrained the selected RNNs to account for monkeys’ irrational choices when making decisions under risk. Importantly, these retrained RNNs eventually make out-of-sample behavioral and neural predictions that generalize across trials and individuals. We also showed that their peculiar wiring induces deterministic interferences in value computations that explain the irrational variability of monkeys’ choices across within-trial attentional trajectories. Finally, we compared the potential biological benefits of rational and irrational variants of OFC circuits and show that the latter exhibits greater tolerance to damage or noise. Irrational interferences in value computation may thus be understood as an incidental byproduct of selective pressure favoring the robustness of OFC circuits to anatomical damage.

That irrational behavior is the incidental outcome of neurobiological constraints is not a novel idea. However, to our knowledge, this work is the first attempt to compare distinct types of constraints and eventually demonstrate the importance of tolerance to circuit damage in this context. We contend that this demonstration is theoretical in essence, at least when compared to empirical work that employ causal – e.g., optogenetic – manipulations to disclose proximal neurobiological constraints^38,39^. Arguably however, it would have been difficult to provide direct empirical evidence for our claim, at least in mammals. This is inherent to the distal nature of the constraint, which is more readily addressed from a computational perspective. In turn, our conclusions rely on a set of modeling assumptions: we will now discuss these.

To begin with, we restricted the set of candidate OFC computations to variants of value synthesis and value comparison. Although a few recent empirical studies consider other types of OFC computations^65^, this prior selection is representative of current debates regarding OFC’s contributions to decision making^1^. Importantly, we showed that some of these variants reproduce complex features of the OFC’s representational geometry, even without being informed with behavioral and/or neural data. This includes established results regarding the mixed selectivity of OFC neural populations (cf. *option value cells, chosen value cells* and *choice cells*)^56,57^. Moreover, we showed that these computational scenarios are anatomically specific, in that their neural predictions do not resemble electrophysiological recordings in either dlPFC or ACC. Retrospectively, this assumption may thus not be so restrictive. Note that the specific RNN variants that we validated using OFC single unit recordings in monkeys are consistent with landmark fMRI studies of value coding in the human vmPFC. In particular, our results directly confirm fMRI studies promoting the attentional format of value coding^23^.They are also consistent with other results; for example, if a default option can be identified prior to decision onset (e.g., in terms of a prior preference over superordinate categories), then pre-stimulus activity in the vmPFC varies with its subjective value, and the magnitude of these variations predicts peoples’ irrational attachment to their default preference^22^. In other words, the vmPFC may use a value coding format that rather distinguishes default versus alternative options. Interestingly, this also aligns with our neural and behavioral results, under the assumption that early preferences – e.g., based upon the first attended cue – set a default option. The reason is twofold. First, as long as attention remains focused on the first option, attentional and default/alternative value-coding formats are formally indistinguishable. Second, the persisting value trace of the firstly attended cue will, on average, appear as a bias towards the default option. In summary, although the statistical resemblance to the default/alternative hypothesis may be stronger in trials where decisions are triggered prematurely – i.e., before all relevant cues have been processed – we argue that our findings remain compatible with known representational frameworks of value coding in the human vmPFC.

Beyond value-coding format issues, one may find it disappointing that we could not disambiguate computational scenarios of value comparison and value synthesis. The underlying question here is whether the OFC directly implements choice, or whether its role is limited to assigning values to available options^49,66^. When implemented in the form of winner-take-all networks, the former scenario explains established findings in electrophysiological and neuroimaging studies, in particular: the observed mixed selectivity of OFC cells^56,60^, as well as the apparent encoding of the value difference between chosen and unchosen options – at least close to choice onset (not shown)^67^. Interestingly, we have shown that such findings can be equally well reproduced by RNNs performing either value synthesis or value comparison, which calls for experiments that are designed to distinguish these kinds of computational scenarios, as opposed to testing one of them.

Also, we did not vary the global architecture of our artificial neural nets, which consisted of a layer of attribute-encoding units sending their outputs to a layer of recurrently connected integration units. In line with recent neural net approaches to value computations in the OFC^20,49^, we adopted the minimal architecture that ensures universal approximation capabilities while using a limited number of artificial units^68,69^. Note that a major computational bottleneck of both value synthesis and value comparison scenarios is OFC circuits’ capacity for combining value-relevant attributes of arbitrary number and type^55^. Now, the above two-layer architecture provides a flexible and simple solution to this problem that rests on the second layer’s trained ability to integrate arbitrary sequences of attributes, whose type and rank are encoded in separate pools of the first layer units. This circumvents the need for otherwise unrealistic, context-dependent changes in connectivity with upstream brain systems involved in recognizing or storing value-relevant information. Nevertheless, the relative simplicity of our design contrasts with previous studies that favored off-the-shelf deep neural nets to approximate the hierarchical organization of, e.g., primates’ visual ventral stream^58^ or humans’ language networks^70^. From a machine learning perspective, tasks such as visual perception and speech comprehension are inherently difficult problems, which remained unsolved until the advent of deep neural nets trained on massive, labeled datasets. In these domains, objective task performance reliably predicts statistical similarity with neural data. This relationship, however, does not readily generalize to our findings: RNNs resemble more closely OFC data when they permit systematic, error-inducing interferences. In retrospect, it is remarkable that our value synthesis/comparison RNNs exhibit such realistic features, at both the behavioral and neural levels. This is despite the degeneracy of RNN wiring profiles with regard to each type of value computation, which we systematically explored by repeating the training process across many random initializations of RNN parameters. Arguably, the ensuing marginalization process renders our results robust to local minima issues. This statistical benefit would have been prohibitively costly to match using deeper neural net architectures.

One might also argue that rational and irrational RNNs may have been compared in an unfair manner. For example, we chose to train rational RNNs under a normative approach, which precludes idiosyncratic variations in risk attitudes. The rationale here was twofold. First, we aimed at selecting neural nets that could serve as neutral and fully interpretable reference points, in that their computational objective was under our control – i.e. computing expected values, as prescribed by rational decision theory. Second, we wanted to assess the neural realism of idealized OFC network models that were derived from first principles alone, without any access to data from the decision task. When evaluating the resemblance of candidate RNNs to OFC recordings, this leaves little room for methodological objections to the ensuing model selection. Now, we acknowledge that, when it comes to measuring statistical similarity to neural recordings, irrational RNNs may benefit from being trained on individual behavioral datasets. However, the fact that irrational RNNs make out-of-sample predictions that generalize across trials and individuals rather suggests that they have captured hidden, yet shared, decision mechanisms. In any case, there is no reason to think that this training difference would favor irrational RNNs in terms of, e.g., tolerance to circuit damage. A related concern is whether the latter may be the artefactual byproduct of re-training *per se*, which may provide an additional opportunity for improving efficiency or robustness. This is the reason why we also explored another training strategy for irrational RNNs, which starts from the same randomly initialized parameter sets as rational RNNs. Reassuringly, our conclusions remain unchanged under this alternative training strategy (see Supplementary Materials).

In conclusion, we believe our modeling assumptions are tenable, at least when compared to state-of-the-art computational studies in the field. Importantly, they have enabled us to reverse the usual approach to disclosing distal neurobiological constraints on rationality, which typically rests on highlighting conflicts with the demands of behavioral performance (cf. Fig. 1c, 1d and 1e). In contrast, we identify realistic mechanisms that explain observed deviations to rationality and explore their potential neurobiological advantages. We believe that this may be a fruitful method for investigating related evolutionary or developmental issues in cognitive neuroscience.

## Methods

### Task design

The decision task is summarized in Fig. 1a. Monkeys were seated in a behavioral chair with their heads restrained. Each trial began when the monkey fixated on a central fixation cue for 500 ms. At the start of the trial, two options were presented, each consisting of two hidden cues initially masked by grey squares. One of these squares then turned blue, indicating the first cue available for sampling. When the subject fixated on the blue square, the corresponding picture cue was revealed and had to be continuously fixated for 300 ms before it was re-masked. All picture cues had been previously learned and were associated with a specific rank of a specific reward attribute (either probability or magnitude). Probability cues indicated reward probabilities of 10%, 30%, 50%, 70%, or 90%, while magnitude cues represented reward magnitudes of 0.15, 0.35, 0.55, 0.75, or 0.95 (arbitrary units of an appetitive juice). Following the initial cue, a second blue square highlighted the next available cue, which had to be sampled using the same procedure. This second cue was either the other attribute of the same option (*option trial*) or the same attribute but for the other option (*attribute trial*). After the second cue, the two remaining cues were simultaneously highlighted with blue squares, allowing the subject to freely choose which one to sample next, or to select one of the two options using a joystick. If a third cue was sampled, the subject could then either sample the final cue or make a choice. Once the fourth cue was revealed, the subject was required to make a choice.

### Value profile estimation

We estimated the subjective value profile of each monkey (and each model) using standard statistical procedures, based on the observed choices (cf. Figures 3a, 3b and 3c). More precisely, we fitted the underlying value function, under the assumption that choices followed a simple softmax mapping of the difference in option values:

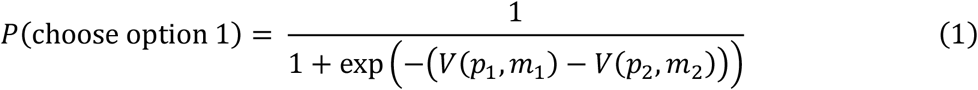

where *p*_*i*_ and *m*_*i*_ denote the reward probability and magnitude of option *i*, as known by the agent at the time of choice, and *V*(*p, m*) is the corresponding subjective value. Equation 1 provides a binomial likelihood function for observed choices, given the unknown monkeys’ value function. Parameterizing the value function then enables us to regress trial-by-trial choices against option attributes.

To maximize modelling flexibility, we employed a semi-parametric approach, whereby each possible combination of probability and magnitude - including cases in which one or both attributes were unknown at the time of choice – is captured using a specific model parameter that quantifies the corresponding value. Given that each attribute has five possible ranks (plus the unknown attribute case), this means that we estimate 6×6=36 parameters from the choices. This semi-parametric approach enables us to capture entirely arbitrary value profiles over the bi-dimensional space spanned by reward probability and magnitude, under the constraints that (i) the same value function applies to all options, and (ii) the value function does not depend upon the cue presentation order. The ensuing probabilistic logistic regression was performed using the VBA academic freeware^71^, which relies on the variational Laplace approach to approximate Bayesian inference^72,73^.

### RNN architecture

The basic RNN architecture is summarized in Fig. 1b. Let *t* ∈ {1,2,3,4} denote the time step at which cues are revealed or attended within a decision trial. The RNN component variables are defined as follows:

- ***U***(*t*) ∈ ℝ^3^: input vector at time *t*, whose entries include the attribute rank (from 1 to 5) and type (probability or magnitude) of the currently attended cue, as well as the identity of the currently attended option (see below).
- ***X***_1_(*t*) ∈ ℝ^9^: unit activation vector in the first hidden layer at time *t*. This *input layer* is com-posed of units that are selective to input entries (units 1 to 5, 6 to 7, and 8 to 9 receive *U*_1_(*t*), *U*_2_(*t*) and *U*_3_(*t*), respectively). It forms a standard population code of the currently attended cue.
- ***X***_2_(*t*) ∈ ℝ^10^: unit activation vector in the second hidden layer at time *t*. Feedforward con-nections convey the current activity of the first layer (***X***_1_(*t*)) to the second layer’s units. But these units are also connected with each other through recurrent connections that carry reentrant dynamics (***X***_2_(*t* − 1)). If properly wired (see below), this *integration layer* can thus integrate currently and previously attended cues.
- ***Y***(*t*) ∈ ℝ^1^ (for value comparison models) or ***Y***(*t*) ∈ ℝ^2^ (for value synthesis models): value readout at time *t*.

At any time *t* within a decision trial, cue-related information propagates through the network according to the following equations:

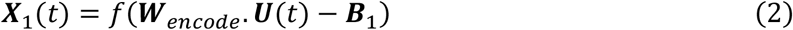

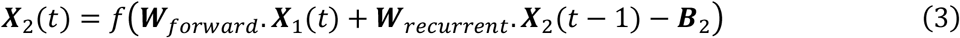

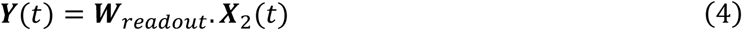

under the constraint that activity within the integration layer is reset at each decision trial, i.e. ***X***_2_(0) = 0 by convention.

Here, ***W***_∎_ refers to matrices of connection weights, and ***B***_1:2_ are bias vectors applied to the corresponding hidden layers. The weights ***W***_*encode*_ and biases ***B***_1_ are set such that each admissible cue rank (*U*_1_) preferentially activates a dedicated unit in a rank-specific pool of first hidden layer units. Similarly, each admissible cue type (*U*_2_) and option identity (*U*_3_) preferentially activates one out of two units each (again in secluded pools of first hidden layer units). To ensure distributed encoding within each pool, the activation profiles of first layer units were set to tile the domain of their specific input uniformly: whenever one unit’s activity reached 75% of its maximum, the next “adjacent” unit in the pool was 25% active.

To impose bounds on units’ firing rates, we use a standard sigmoid activation function *f* for all hidden units:

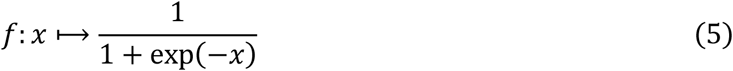

Importantly, when structurally organized into two such hidden layers, neural nets with a limited number of sigmoidal units possess universal approximation capabilities^68,69^. This means that RNNs of this type can be trained to store memory traces of all previously attended cues, albeit in a format that may not be directly accessible. More generally, it should be possible to train such RNNs (see below) to compute any arbitrary mapping of the sequence of attended cues.

The RNN receives inputs one at a time, in a sequential manner, as monkeys did in the task. The sequence order is determined by the exogenous control of attention, which samples cues in an arbitrary fashion within a decision trial. Let *U*_1_(*t*), *U*_2_(*t*) and *U*_3_(*t*) denote the entries of the input vector ***U***(*t*) ∈ ℝ^3^:

- *U*_1_(*t*) encodes the normalized rank of the attended cue, with the following mapping:

| Magnitude cue | Probability cue | Cue rank | $U_1$ |
| --- | --- | --- | --- |
| 0.15 AU | 10 % | 1 | 0.1 |
| 0.35 AU | 30 % | 2 | 0.3 |
| 0.55 AU | 50 % | 3 | 0.5 |
| 0.75 AU | 70 % | 4 | 0.7 |
| 0.95 AU | 90 % | 5 | 0.9 |

- *U*_2_(*t*) encodes the attribute type, i.e. probability: *U*_2_ = 0 and magnitude: *U*_2_ = 1.
- *U*_3_(*t*) encodes the identity of the attended option, i.e. option 1: *U*_3_ = 0, option 2: *U*_3_ = 1.

Note that the identity of the attended option can be encoded in two different representation formats: spatial (left vs. right) or temporal (first vs. second). This distinction affects the encoding of *U*_3_, as illustrated in the following example trials:

| Trial ID | Time step | Attended option side | $U_3$ in the *spatial* frame<br>(right = 0, left = 1) | $U_3$ in the *temporal* frame<br>(first = 0, second = 1) |
| --- | --- | --- | --- | --- |
| 1 | 1 | Right | 0 | 0 |
| 1 | 2 | Left | 1 | 1 |
| 1 | 3 | Left | 1 | 1 |
| 1 | 4 | Right | 0 | 0 |
| 2 | 1 | Left | 1 | 0 |
| 2 | 2 | Left | 1 | 0 |
| 2 | 3 | Right | 0 | 1 |
| 2 | 4 | Right | 0 | 1 |

Although the encoding frame of the attended option’s identity does not modify the raw information that is conveyed to the network, it nonetheless determines the pattern of activity in the input layer. In turn, the encoding format is likely to modify the sequence of activity patterns of the whole RNN over time steps within a decision trial. In other terms, the way the RNN responds to a sequence of cues will depend upon the encoding format of the attended option’s identity.

Similarly, the value outputs ***Y***(*t*) of the network can be expressed in different representation formats: spatial, temporal, or attentional (attended vs. unattended). The example trials below illustrate how the encoding format of option values varies across these frames. Let *V*_*left*_ and *V*_*right*_ denote the values of the left and right options as estimated by the network at each cue onset. The statistical similarity between representation formats depends on the actual sequence order of cue attendance:

| Trial ID | Time step | Attended option side | $Y$ in the *spatial* frame | $Y$ in the *temporal* frame | $Y$ in the *attentional* frame |
| --- | --- | --- | --- | --- | --- |
| 1 | 1 | Right | $V_{right} \& V_{left}$ | $V_{right} \& V_{left}$ | $V_{right} \& V_{left}$ |
| 1 | 2 | Left | $V_{right} \& V_{left}$ | $V_{right} \& V_{left}$ | $V_{left} \& V_{right}$ |
| 1 | 3 | Left | $V_{right} \& V_{left}$ | $V_{right} \& V_{left}$ | $V_{left} \& V_{right}$ |
| 1 | 4 | Right | $V_{right} \& V_{left}$ | $V_{right} \& V_{left}$ | $V_{right} \& V_{left}$ |
| 2 | 1 | Left | $V_{right} \& V_{left}$ | $V_{left} \& V_{right}$ | $V_{left} \& V_{right}$ |
| 2 | 2 | Left | $V_{right} \& V_{left}$ | $V_{left} \& V_{right}$ | $V_{left} \& V_{right}$ |
| 2 | 3 | Right | $V_{right} \& V_{left}$ | $V_{left} \& V_{right}$ | $V_{right} \& V_{left}$ |
| 2 | 4 | Right | $V_{right} \& V_{left}$ | $V_{left} \& V_{right}$ | $V_{right} \& V_{left}$ |

Note that not all combinations of input/output formats are admissible. More precisely, when the currently attended option’s identity is encoded using the spatial format, then value outputs can be encoded in all representation formats (3 possibilities). However, when the currently attended option’s identity is encoded using the temporal format, then the spatial information is lost, which leaves only 2 possible value encoding formats (temporal and attentional frames). This also means that there are only 5 combinations of input/output representation formats in total. Training the RNN to encode value computations within a given format will require specific settings of recurrent connection weights. This implies that the encoding format of value readouts is also likely to modify the way the RNN responds to a sequence of cues.

### RNN training

#### Rational training

Under standard decision theory, rational choices maximize the *expected value*, where the expected value of an option is defined as the product between reward probability and magnitude: *EV* = *p* × *m*. Formally, options’ expected value can be defined at each time step within a decision trial, under the assumption that option attributes that are unknown (i.e., yet to be sampled) can be replaced by their expectation under the task distribution. In principle, *value synthesis* and *value comparison* models can thus be trained to output the expected value of both options or, respectively, the difference in expected values, in response to each cue presentation (at each time step within a decision trial). It turns out that RNNs of the sort we describe above can reliably learn to solve the problem of integrating previously and currently attended cues to output options’ expected values from almost any random initialization of their connectivity.

All RNN models were implemented and trained using the VBA academic freeware^71^. The RNN parameters subject to training (***W***_*forward*_, ***W***_*recurrent*_, ***W***_*readout*_ and ***B***_2_) were initialized as samples from an i.i.d. Gaussian distribution with mean 0 and variance 0.5. For each RNN model, the training procedure was repeated with a different initial random sample, until 1,000 trained models reached 95% accuracy on held-out test data. This enables us to sample the unknown manifold of RNN wirings that can perform a given type of value computation. In the main text, we refer to the ensemble of trained RNNs as a *cohort*, each of which corresponds to a given type of value computation (value synthesis versus value comparison) and a given combination of input/output representation format (see above).

For each model instance in a given RNN cohort, a training set and a testing set consisting of 500 trials each were generated. Every trial consisted of a sequence of four cues, randomly chosen among the set of different option pairings, and presented in a random sequence order. Importantly, trained models carry no memory trace of preceding decision trials (their internal state is reset to baseline levels at each trial onset). Also, we did not endow RNNs with the capacity to decide which cue to attend to or when to commit to a decision; rather, we trained them to operate value computations independently of such processes, which are treated as exogenous and arbitrary. Note that training and testing trials could be classified post-hoc as either *attribute trials* or *option trials*, depending on whether attention switched to the second option at the second cue onset, or not.

Training was terminated when the absolute change in variational free energy between VBA successive iterations fell below 10. However, the trained RNN was kept in its corresponding model cohort only if it reached at least 95% of explained variance (in trial-by-trial expected values) on its testing set. Each RNN cohort consisted of 1,000 independently trained model instances, each with a unique training set, testing set, and parameter initialization. Importantly, random seeds were shared across cohorts, which allowed for paired comparisons across cohorts.

Importantly, we stored the updated model parameters at each step of the training procedure, for each model instance of each RNN cohort. This enabled us to test RNNs’ neural predictions against OFC neural data (see below) as training unfolds (cf. Fig. 2c).

#### Rational training with biological constraints

Decision systems in the brain may have been shaped under the multiple imperatives of producing rational choices and satisfying biological (energetic, informational or robustness) constraints. Under a neural net perspective, all behavioral and biological properties of such systems are determined by their internal wiring. Therefore, training RNNs to perform rational value computations while simultaneously complying with these constraints should enable us to reveal possible conflicts between behavioral and biological imperatives.

Here, we focused on three main biological constraints: namely, maximal information transfer rate, minimal energy budget, and maximal tolerance to damage (see section *Biological constraints* below). In brief, RNN training followed the exact same procedure as above, except that the objective function included an additional biological constraint adequacy term. To control the balance between behavioral and biological imperatives, we introduced a constraint compliance weight hyperparameter that varied exponentially from 10^-3^ to 10^3^ (the higher the constraint weight, the tighter the biological constraint on the RNN wiring) and trained 10 RNN instances per constraint weight. This enabled us to compare RNN cohorts trained under systematically varied levels of biological pressure.

For example, Fig. 1c, 1d and 1e plot the mean rate of rational choices against the achieved biological constraint adequacy level, for each compliance weight (and each biological constraint). This reveals the shape of the ensuing Pareto front, which tells how conflicting behavioral and biological imperatives are. If the biological constraint spans the null space of value computations, then increasing the compliance weight has no bearing on choice rationality. However, if behavioral and biological imperatives conflict with each other, tightening the biological constraint results in larger rationality losses, which is what we see here.

#### Re-training RNNs to explain monkeys’ irrational choices

To begin with, recall that monkeys often commit to a choice before having sampled all cues. However, this does not imply that such choices are irrational, in that monkeys may still select the option with the highest expected value (where unknown attributes are replaced by their expectation under the task distribution). In fact, according to this criterion, only about 20% of monkeys’ choices are irrational. To preserve the interpretability of value computations and input/output representation formats of rational RNNs, the re-training of RNN models was restricted to ***W***_*recurrent*_ (i.e. all other parameters were frozen). This effectively restricts the admissible sources of irrational choices to within-trial nonlinear interferences and spill-over effects between attended cues.

In contrast to the rational training phase, where value outputs can be evaluated at each cue onset within decision trials, irrational re-training relies solely on observed monkey choices. The latter only constrain the value readouts of retrained RNNs at the time of choice (i.e. after the last sampled cue). Moreover, the only available information is the choice itself, which we compare to RNN value readouts via the same softmax mapping as in Equation 1. From this perspective, the retraining of RNNs may be seen as a way to relax the two constraints of the logistic regression in Equation 1 (value invariance across options and cue presentation orders).

Each RNN instance within each cohort was then re-trained to fit the choices of each individual monkey, using a training dataset of 2,000 trials randomly selected from monkeys’ recorded sessions. This procedure produces two twin versions of each retrained irrational model - one for each monkey. We then test their respective behavioral (Figure 3e) and neural (Figure 3f, 3g and 3h) predictions within and across monkeys. This enables us to evaluate their inter-trial and inter-individual generalization ability.

Note that neither rational training, nor irrational retraining is informed by neural recordings. This greatly simplifies the validation of quantitative model predictions regarding the representational geometry of OFC neural populations.

### Analysis of representational geometry within neural populations

#### Representational similarity analysis

Let 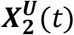 denote the vector of activations in the RNN’s integration layer at time *t* in response to a sequence of input ***U***(1), …, ***U***(*t*). At this first cue onset (*t* = 1), this vector can be computed for each possible input ***U***_*k*_, which yields 20 distinct activation patterns (i.e., 5 cue ranks × 2 cue types × 2 options). The ensuing representational dissimilarity matrix (*RDM*) is constructed element by element by computing pairwise similarities between these activation patterns^74^:

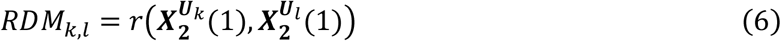

where *r* denotes Pearson’s correlation. If *RDM*_*k,l*_ is strongly positive, then activity patterns are mostly invariant to differences between inputs ***U***_*k*_ and ***U***_*l*_, i.e. the neural representation of these inputs are similar. In brief, RDMs enables us to identify what input features need to change to elicit distinct neural responses.

The same procedure is applied to recordings of OFC neurons (as well as to neural recordings within the dlPFC and the ACC), where 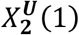 is replaced with the average firing rate in a 100-400 ms window after the first cue onset. This yields two RDMs: one for the model (*RDM*^*model*^) and one for the OFC data (*RDM*^*OFC*^). Full RDM summary statistics for all monkeys and brain regions can be eyeballed in Figure S2.

Finally, the similarity between these matrices is quantified using a rank-based distance metric:

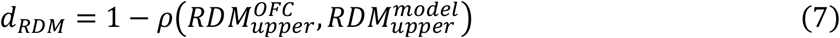

Here, *ρ* denotes Spearman’s correlation and *RDM*_*upper*_ refers to the upper triangular half of the matrix, excluding the diagonal. We used a rank-based metric because experimental neural data is typically much noisier than RNN activations, resulting in compressed correlation ranges that are more appropriately captured by rank correlations.

#### Cross-correlation matrices

The benefit of the above representational similarity analysis is to account for all attributes of a given cue, i.e. attended option identity, attended cue type and attended cue rank. Unfortunately, it does not scale well with the number of cue combinations. In our context, its statistical cost is prohibitive for later phases of decision trials, when more than one cue has been attended. For example, at the second cue onset, there are 20×20=400 possible cue combinations, which would induce RDMs with almost 79,800 elements. This is incompatible with the number of acquired decision trials in the task. Thus, we resort to another type of summary statistics, which was proposed by Hunt and colleagues^50^. In brief, this analysis enables us to quantify and compare the multiple traces that cue sequences leave on units’ activity, at the cost of neglecting differences induced by cue types. This simplifying assumption exploits the observed quasi-symmetrical impact of reward probability and magnitude on monkeys’ subjective value profiles (see Fig. 2a).

Let 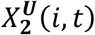 denote the activation of unit *i* in the RNN’s integration layer at time *t* in response to a sequence of input ***U***(1), …, ***U***(*t*). We regress each unit’s trial-by-trial activity variations at cue onset *t* concurrently onto trial-by-trial variations of normalized attribute rank in all cues, while identifying cues by their appearance order in the sequence:

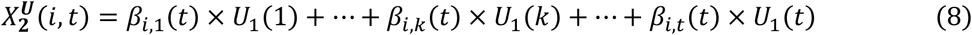

Note that we also include two additional regressors, which encode how consistent the 2^nd^ and 3^rd^ cues are with regard to the currently preferred option, as well as an intercept term (not shown in Equation 6). This approach aims at detecting nontrivial memory traces of previously attended cues, while ruling out mere confirmation effects in value coding neurons. Importantly, we separate *option trials* (where the first two cues belong to the same option) from *attribute trials* (where the first two cues describe the same attribute – i.e. probability or magnitude – but for both options) prior to performing the regression analyses. This yields one set of regression coefficient estimates *β*_*i,k*_(*t*) per trial type, for each unit and each cue in the sequence.

Let 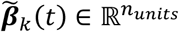 denote the vector of t-statistics associated with regression coefficient estimates for the *k*^th^ attended cue (*k* ∈ {1, 2, 3}), across integration units. This vector measures how sensitive to the *k*^th^ attended cue second layer units are (at time *t*) in normalized signal-to-noise ratio units. This enables a direct quantitative comparison across units, cue presentation orders and decision times. Note that 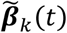 vectors that involve cues presented after the time when units’ activity is sampled (i.e. when *k* > *t*) are statistically meaningless.

We then define the cross-correlation matrix (*CCM*) as follows:

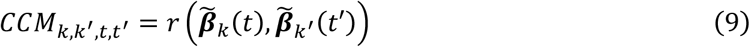

where *r* denotes Pearson’s correlation. A strongly positive CCM cell indicates that those units that are most sensitive to the rank of the *k*^th^ attended cue at time *t* are also those most sensitive to the rank of the *k*′^th^ cue at time *t*′. Importantly, the sign of regression coefficients matters. This means that CCM analyses can detect when a unit respond positively to some cue while responding negatively to another one (as may be the case when cues belong to distinct options).

We obtain full CCMs by systematically varying cue presentation orders (*k* and *k*′) as well as activity sampling times (*t* and *t*′), yielding a 9 by 9 symmetrical matrix. We then remove CCM cells that are meaningless (*t* < *k* or *t*^′^ < *k*^′^) to avoid statistical illusions possibly induced by imperfections in trial randomizations. We repeat this process for both *option trials* and *attribute trials*, yielding two CCM types. Differences between the two types of CCM cells that involve the first and second cue onset times (i.e. *CCM*_1,2,∎,∎_) signal a shift in how the attended option affects the network’s distributed computations. In particular, if neurons respond to the value difference between options, then one expects *CCM*_1,2,2,2_ to be positive for option trials, and negative for attribute trials^50^.

We apply the exact same analysis on recorded data from OFC neurons (as well as neurons in the dlPFC and ACC), where 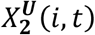 is replaced with the average firing rate in a 100-400 ms window after each cue onset. This provides summary statistics whose temporal resolution matches that of RNN models. The procedure is summarized on Figure M1 above.

**Fig. M1.**
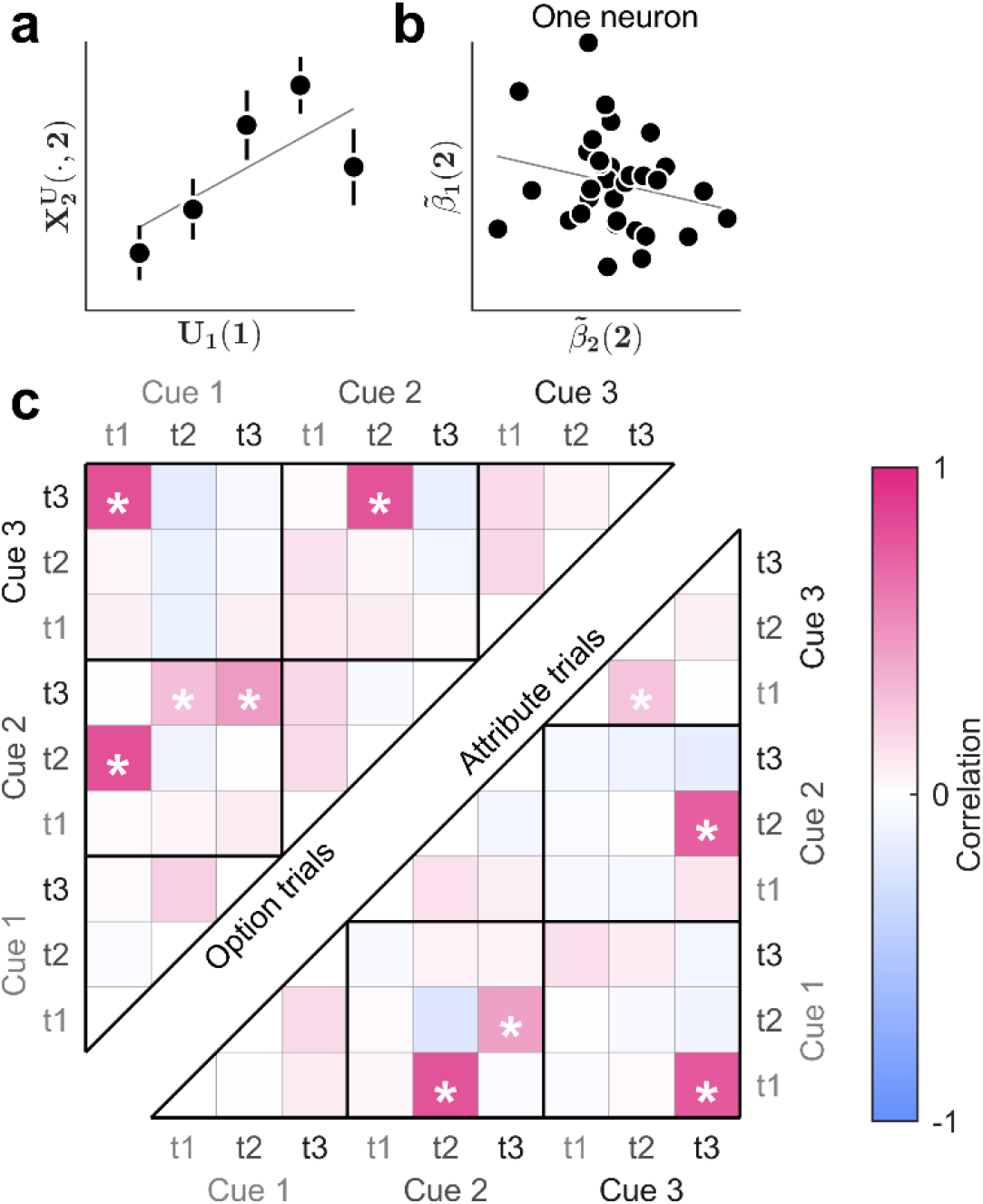
Derivation of CCM statistics in OFC neurons. **a**, First, we regress the mean firing rate of each neuron at each cue onset against the rank of all previously attended cues (across trials). Here, we show the apparent statistical relationship between the activity 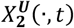 of one example OFC neuron sampled at time *t* = 2 (y-axis), plotted against the rank *U*_1_(*k*) of the first presented cue (*k* = 1). **b**, Second, we measure the correlation (across neurons), between the ensuing regression coefficients for different activity sampling and cue presentation times. Here, we plot the sensitivity 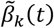 of OFC neurons (one dot is one neuron) sampled at time *t* = 2 to the first cue (*k* = 1, y-axis) against their sensitivity to the second cue (*k* = 2, x-axis). **c**, Each cell in the CCM matrix shows the correlation across neurons for a given pair of regression coefficients (neurons pooled across monkeys). The upper half of the matrix shows the results computed on *option trials* (where the two first cues characterize the same option), while the lower half corresponds to *attribute trials* (where the two first cues characterize the same attribute, but different options). Asterisks indicate significant correlations, with p-value < 0.0007 (correction for multiple comparisons across CCM cells).

To compare the informational geometry of RNNs and OFC neural populations, we simply compute the Euclidian distance between the meaningful CCM cells:

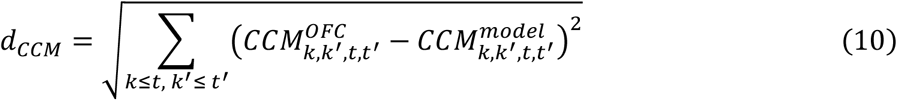

Note that, although this approach scales well with the number of cue combinations, it typically is insensitive to the type of attribute (probability vs magnitude) of attended cues.

#### Mixed selectivity: offer value cells, chosen value cells and choice cells

To identify possible “offer value”, “chosen value”, and “choice” cells, we replicated the analysis previously introduced by Padoa-Schioppa and colleagues^57^. For each unit, we performed four separate regressions of activity sampled at the time of choice, across all trials, against the value of option 1, the value of option 2, the value of the chosen option (defined as the monkey’s recorded choice), or the identity of the chosen option, respectively. Each unit was assigned to the category that yielded the highest percentage of explained variance, provided the regression was significant (p-value < 0.05). Otherwise, no category was assigned. When applied to neural recordings in the OFC, we relied on subjective value profiles, as estimated from monkeys’ choices in the task (see subsection Value profile estimation above). To maximize the match between analyses, we also use model-specific value profiles for RNNs (using the encoding formats that corresponds to each RNN model).

### Analysis of computational interferences in irrational RNNs

#### Dependency on cue sequence order

In principle, rational behavior in the task only depends upon the content of value-relevant information, but not on its presentation sequence order. Under this view, any observed dependency on cue sequence order violates rationality.

Let Δ*V*^***U***^(*t*) denote the value difference between options, as can be readout from the RNN’s activity pattern at time *t*, in response to a given sequence of input ***U***(1), …, ***U***(*t*). Note that, for value synthesis models, we compute Δ*V*^***U***^(*t*) by simply subtracting the readouts of both option values. To quantify the dependency on cue presentation order, we first simulate the RNN response to all admissible permutations of cue orderings while keeping the combination of *t* attended cues constant, measure the standard deviation of Δ*V*^***U***^(*t*), and then average the results over all possible cue combinations. Importantly, we repeat this process separately for option trials and attribute trials, meaning that we only consider cue order permutations that are admissible for each trial type.

Let 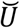 be the set of all possible combinations of *t* cues, and for each member set 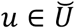, let *S*(*u*) denote the set of admissible permuted sequence orderings of those cues (restricted to the relevant trial type). Then, we define the dependency on sequence order at time *t*, denoted *d*_*S*_(*t*), as follows:

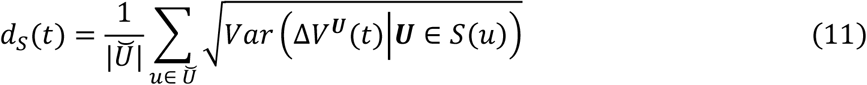

Note that *d*_*S*_(*t*) is defined for all decision times starting from the second cue onset up until the last cue onset (2 ≤ *t* ≤ 4). As we show in the Results section, eyeballing *d*_*S*_(*t*) as a function of decision time enables us to track the possible accumulation of interferences in RNN computations. Reassuringly, rational RNNs show no detectable dependency on sequence order, i.e. *d*_*S*_(*t*) ≈ 0, irrespective of cue onset time (not shown).

Note that this analysis cannot be directly applied to monkeys’ choices, as we cannot have access to the monkeys’ internal value estimates for each cue sequence order. This is because the total number of admissible cue sequence orders is prohibitive, at least for late cue inset times. This means that decision trials in the task do not sample the set of admissible cue sequence order in a dense manner. Therefore, we resort to simpler measures of apparent deviations to rational choice, which effectively reduce to detecting trials that are incongruent with monkeys’ value profile estimates (see subsection Value profile estimation above). This effectively yields a trial-by-trial binary rationality flag: its sample average provides the apparent rate of rational choices for each decision time (2 ≤ *t* ≤ 4). Now, monkeys’ choice stochasticity may render difficult decisions more likely to violate strict preference orderings. We thus need to correct the apparent rate of rotational choices for decision difficulty. To do this, we perform a logistic regression of the trial-by-trial rationality flag variable onto decision difficulty, as given by the absolute difference in estimated subjective option values (separately for each monkey). The residuals of this regression quantify the amount of trial-by-trial rationality that cannot be explained by variations in decision difficulty. The sample average of these residuals, for each decision time, is what we call the corrected rate of rational choices.

#### Persisting value traces

The above dependency on sequence order may be partly driven by a directional bias, whereby the effective weight of each cue is determined by its onset time. For example, previously attended cues may weigh more on value outputs than currently attended cues, all else being equal. We developed a specific method for detecting such persisting value traces, which can be equally applied to both RNN simulations and monkeys’ behavior in the task.

We start by re-estimating value profiles, while allowing for value differences between options that are currently or previously attended (at the time of choice), and having separated trials by the type of attended cue (reward probability vs magnitude). Let 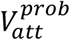 denote the pseudo-value function of the attended option when a probability cue is attended at the time of choice, and 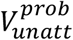 that of the other (unattended) option. Let *p*_*att*_ and *m*_*att*_ be the ranks of the attended option’s probability and magnitude, and *p*_*unatt*_ and *m*_*unatt*_ those of the unattended option. The probability of choosing the attended option is given by:

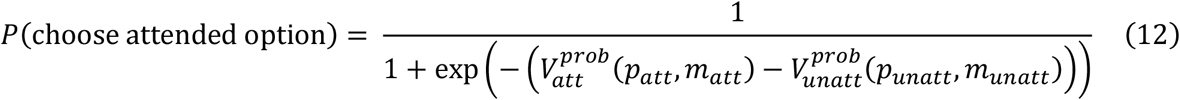

This provides a binomial likelihood function for observed choices that are triggered when a probability cue is attended. To estimate the pseudo-value profiles 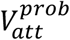 and 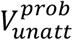, we use the same semi-parametric approach as before (see subsection Value profile estimation above). The pseudo-value profiles 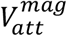 and 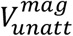 can be estimated similarly, given observed choices that are triggered when a magnitude cue is attended.

Recall that 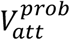 (resp., 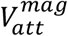) is the pseudo-value that ensues from currently attending a probability (resp., a magnitude) cue, while the magnitude (resp., probability) cue was previously attended (if ever). To quantify the relative impact of currently and previously attended cues while marginalizing over cue types, we then combine 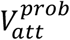 and 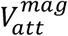 to form the following average pseudo-value profile *V*_*att*_:

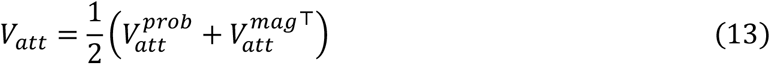

Importantly, *V*_*att*_ is a 6×6 pseudo-value profile whose first dimension (columns) spans the rank of the currently attended cue, while its second dimension (rows) spans the rank of the previously attended cue – including the case where it is unknown at the time of choice. Note that rational agents exhibit a strictly symmetric average pseudo-value profile, irrespective of their subjective value profile over the bidimensional space spanned by native option attributes. However, irrational value computations that induce persisting value traces do exhibit asymmetrical *V*_*att*_ profiles.

To quantify potential asymmetries in *V*_*att*_, we regressed the ranks of the attended and unattended cues onto *V*_*att*_ using a generalized linear model:

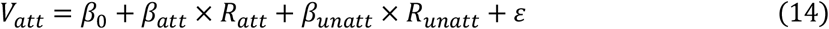

where *R*_*att*_ and *R*_*unat*_ denote the ranks of the attended and unattended cue, respectively, each ranging from 1 to 5, whereas *β*_*att*_ and *β*_*unatt*_ quantify the contribution of attended and unattended cues to the pseudo-value profile. In the Results section, we report the difference Δ*β* = *β*_*att*_ − *β*_*unatt*_ between estimates of attended and unattended cue contributions. If Δβ < 0, then the pseudo-value profile *V*_*att*_ is asymmetric and more sensitive to the unattended cue. This is the hallmark of a persisting value trace that resists novel (currently attended) information.

### Biological constraints

We now derive measures of biological constraint compliance, as can be derived from extensive numerical simulations of trained (rational or irrational) RNN models of the OFC. In what follows, *N* = 10, |*U*| = 10,000 and *T* = 4 are the number of units in the integration layer, the number of simulated input sequences and the number of time steps within a given decision trial.

#### Energetic budget: average network firing rate

The brain’s energetic budget constraints are tight. Since action potentials are a major source of energetic consumption in neurons, we quantified the pseudo-energetic budget *Ē* of an RNN in terms of the average activation of RNNs’ integration units, across all units, time steps, and admissible cue sequences:

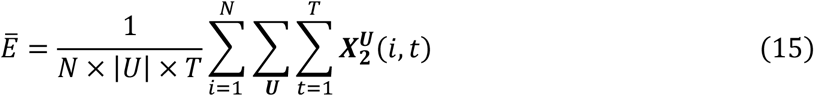

Where 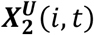 is the activity level of unit *i* at time *t* in response to the input sequence ***U***. Since units’ activation functions are standard sigmoid mappings, 0 ≤ *Ē* ≤ 1.

#### Efficient coding: code redundancy and information transfer rate

Efficient coding models suggest that brain circuits with limited neural resources self-organize to either minimize code redundancy or maximize information transfer rate.

First, recall that population codes are redundant if neural elements tend to be active at the same time, across the range of stimuli that drive their responses. Here, a unit *i* is deemed “active” if its output ***X***_2_(*i*) exceeds the *a*^th^ percentile of its marginal activity distribution (see below). Let *N*_*active*_(*a*, ***U***, *t*) denote the number of active units at decision time *t*, for the input sequence ***U***, under the exceedance threshold *a*. The probability that two randomly selected units are simultaneously active is given by:

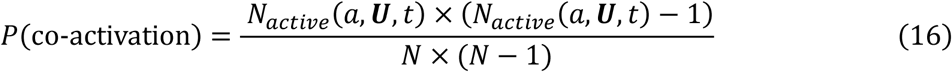

The higher the co-activation probability, the more redundant the neural code. However, changing the exceedance thresholds modifies the co-activation probability estimate. In analogy to Receiver Operating Characteristic (ROC) analyses, we thus systematically vary the exceedance threshold *a* ∈ {0, 1, …, 100}, and derive the ensuing probability of co-activation. We then define the code redundancy 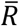 as the average probability of co-activation over exceedance thresholds:

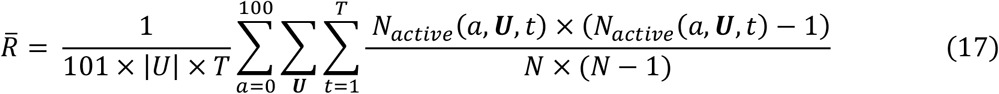

By construction, 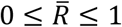. When 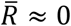, code redundancy is minimal, i.e. units almost never co-activate across trials and decision time steps.

Second, we measure information transfer rate in terms of the average entropy of the unit’s output (across integration units). Let *f*: *x* ↦ *y* be the input-output activation function of neural net units. At the low noise limit, information transfer rate *Ī* reduces to the expected, log-transformed, absolute gradient of the activation function^64^:

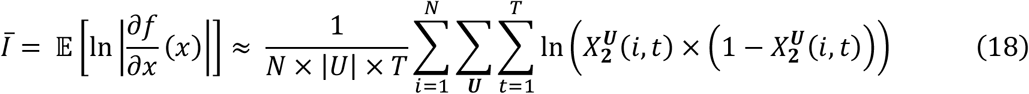

where the expectation is taken under the distribution of admissible cue sequences ***U*** and we have used standard results of sigmoidal activation functions. Note that the maximum information transfer rate is achieved when the distribution of inputs to each integration units exactly matches the gradient of their activation function (here: *Ī ≲* −1.38 nats).

#### Robustness: excitatory/Inhibitory balance and tolerance to damage

In this work, we consider two distinct types of robustness.

First, the excitatory/inhibitory balance of a circuit is critical for maintaining its dynamical stability. Formally, it refers to the relative contribution of excitatory and inhibitory inputs on features of the circuit’s evoked responses (e.g., selective tuning). In electrophysiological studies, E/I balance is typically evaluated using intracellular conductance estimates across a wide range of conditions and contexts. Here, we quantify a simple form of structural balance 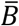, which we define as the ratio between the number of positive (excitatory) and negative (negative) connection weights:

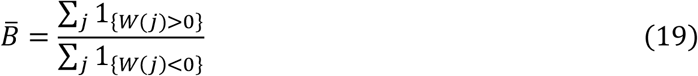

where 1_{*W*(*j*)≶0}_ is binary indicator that flags whenever the *j*^th^ entry of the augmented connection weight matrix ***W*** = ***W***_*forward*_ ⋃ ***W***_*recurrent*_ is positive or negative. By construction, 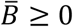. Note that we expect rational RNNs to be relatively well balanced 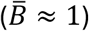, because excessive excitatory or inhibitory connections may induce runaway or saturating activity dynamics, which precludes accurate value computations (at least in late phases of decision trials).

Second, biological systems need to maintain function despite compromised structural integrity. Here, we define the RNN’s tolerance to damage in terms of their achieved rate of rational choices under varying levels of artificial lesions on the integration layer. Let *n* ∈ {1,2, …, 9} and ***C***_*n*_ denote the number of lesioned units in the integration layer, and the *N* × 1 binary lesion map vector that indicates the combination of lesioned units, respectively. Artificial lesions are performed by forcing lesioned units to stay silent across all time steps and trials. Let *z*_*model*_(***U***, *t*, ***C***_*n*_) ∈ {0, 1} denote the RNN’s simulated choice at time *t* in response to an input sequence ***U***, under a lesion ***C***_*n*_ of its integration layer. Let *z*_*rational*_(***U***, *t*) denote the rational choice (i.e., the preferred option based upon options’ expected value) given the same input sequence. We define the tolerance to damage 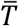 as the retained rate of rational choice, averaged over all possible lesion maps involving 10% to 50% of all units in the integration layer:

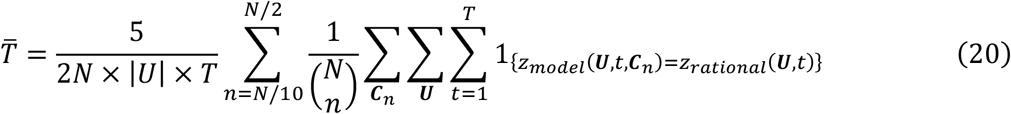

where 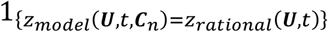 is a binary indicator that flags whenever the lesioned RNN emits a rational choice, and 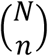 is the number of possible combinations (lesion maps) when lesioning *n* among *N*. By construction, 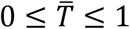. If 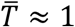, then RNNs’ value computations are mostly unaffected by virtual lesions. We note that the results we report in the article are mostly invariant to the chosen range of lesion extent (here: from 10% to 50%).

