## Supplemental Materials for "Irrational decisions reflect robustness constraints on value computations implemented by orbitofrontal circuits"

**SUPPLEMENTARY MATERIALS**

### 1. How specific is the information content of rational RNNs?

In the main text, we evaluate the resemblance of rational RNNs to OFC neurons' activity patterns, under the assumption that each RNN cohort entertains a specific type of internal representations (despite yielding identical decisions in the task). To provide evidence for this assumption, we performed the following decoding analysis. First, we evaluate each RNN instance of each cohort on the 500 decision trials they were trained on. For each simulated set of activity patterns, we train a linear decoder of option values in all output readout formats, separately in terms of the pair of option values, or in terms of the value difference between options. Finally, we test the linear decoder on a set of 500 different decision trials. We then measure the decoding accuracy on the test dataset using the relative predicted residual error sum of squares (rPRESS) statistic:

$$rPRESS = \frac{\sum_{t=1}^{T}(y_t - \hat{y}_t)^2}{\sum_{t=1}^{T}\left(y_t - \frac{1}{T}\sum_{t=1}^{T} y_t\right)^2}$$

where $y$ and $\hat{y}$ are the actual and predicted option values, respectively (and $T$ is the total number of simulated epochs).

We then average the rPRESS statistics over instances within each RNN cohort. Figure S1 below summarizes the results of this analysis.

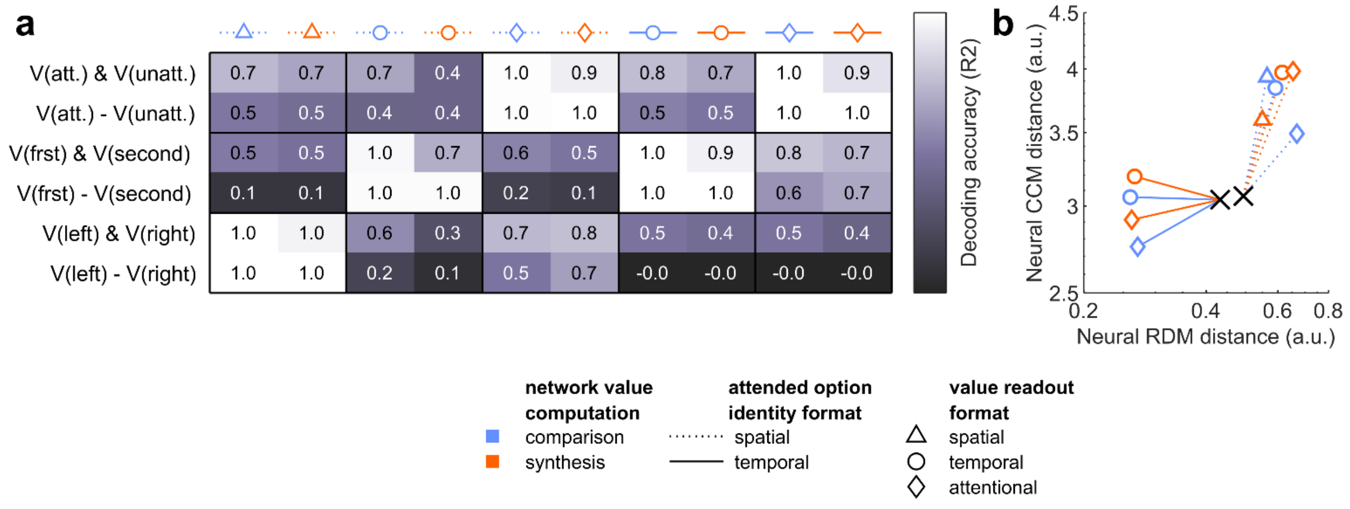


**Fig. S1 | Specificity of information content in RNN cohorts. a**, Decoding accuracy of each type of output (lines) is shown for each RNN cohort (columns). **b**, Neural distance trajectories between OFC and RNN cohorts during rational training. Points show the average distance between activity patterns of OFC recordings and RNN models (across the 1,000 RNN instances), computed using either RDMs (x-axis) or CCMs (y-axis) metrics, either at initialization (black crosses) or at training convergence (see legend). This is a simple summary of Fig. 2c in the main text.

In brief, the specific value computation that is performed by the RNN matters. More precisely, it is almost impossible to reliably decode option values framed in a given option identity format from response patterns of RNNs that were trained under different option identity formats. Also, RNNs that perform value comparison have lost some information about the pair of option values. This is evident when comparing the decoding accuracy for pairs of option values in either value comparison or value synthesis RNNs.

### 2. How different are the representational geometries of the OFC, dlPFC and ACC?

In the main text, we compare RNN cohorts in terms of their representational geometry. We do this w.r.t two distance metrics, based on either RDMs or CCMs, respectively. Recall that RDMs quantify how dissimilar evoked activity patterns are for any pair of cue (2x2x5 = 20 possibilities at first cue onset). In contrast, CCMs quantify the similarity of profiles of neural sensitivity to present and past cue ranks. Although CCM-based distance metrics are insensitive to cue type (probability or magnitude), they quantify potential internal memory traces about previously sampled cues. Therefore, one can think of RDM- and CCM-based distances as two complementary metrics of similarity between RNN models and neural populations.

First, Figure S2 shows the raw RDM matrices of each monkey and pooled recorded neurons of both monkeys together, in either the OFC, dlPFC or ACC.

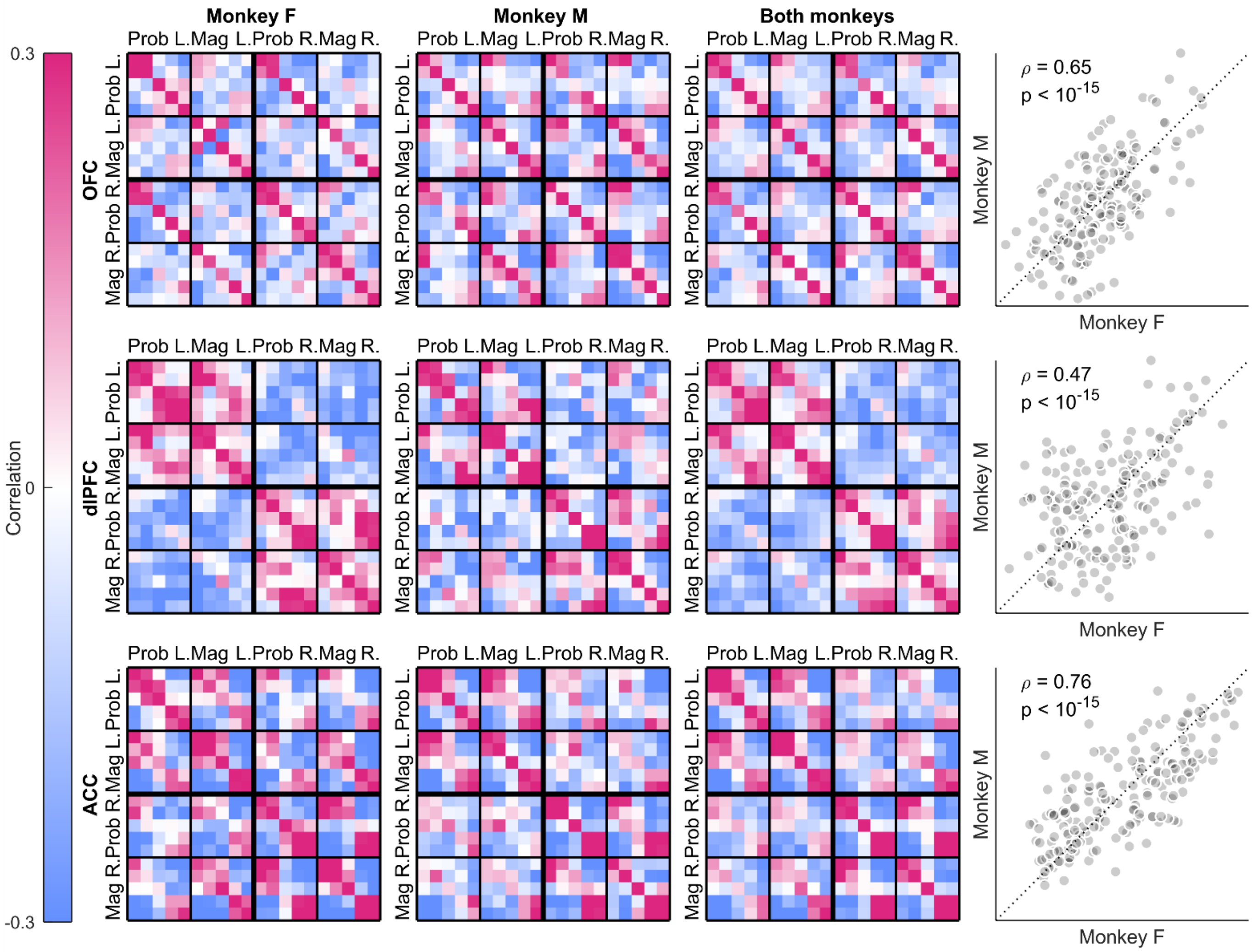


**Fig. S2 | RDMs at first cue onset for neural populations recorded in either the OFC, dlPFC or ACC.** Panels on the left show RDMs (color code depicts the Pearson correlation coefficients between activity patterns that are elicited in response to first cue, going from blue – low correlation – to pink – high correlation –), organized by cue spatial side (left: L or right: R), type (probability or magnitude) and rank (from 1 to 5). Panels on the right show RDM cells from monkey M (y-axis) plotted against RDM cells from monkey F (x-axis).

The representational geometry of OFC, dlPFC and ACC seems to be regionally specific. For example, both dlPFC and ACC neurons seem to respond differently when cues are presented on either the left or the right side (cf. dampening of neural similarity in extra-diagonal blocks of RDMs). This is not the case for OFC neurons, which exhibit similar responses to cues presented on either the left or right side (cf. high extra-diagonal RDM entries). In addition, dlPFC and ACC neurons seem to respond very similarly to probability or magnitude cues, which is less evident in OFC neurons.

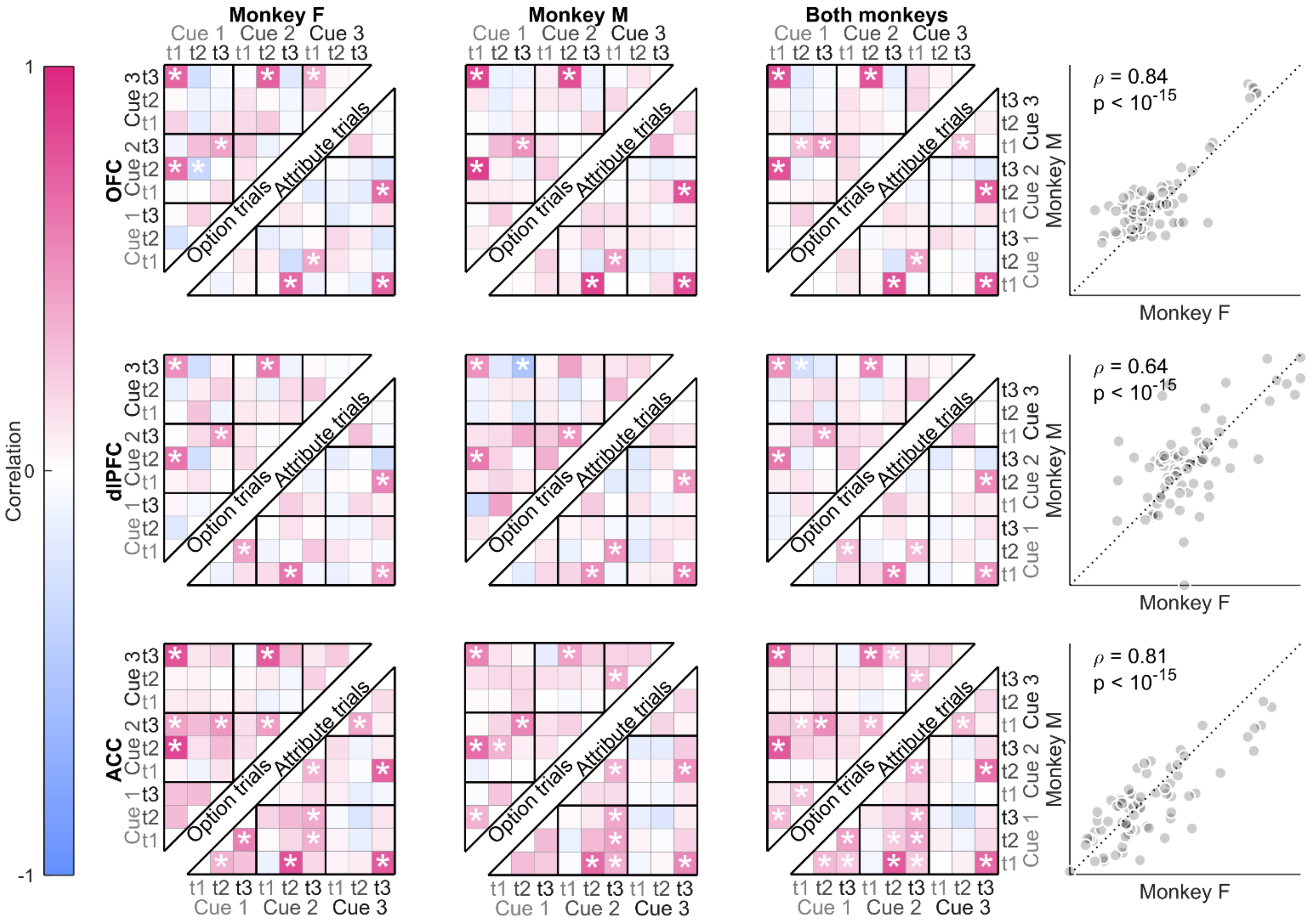


**Fig. S3 | CCMs for neural populations recorded in either the OFC, dlPFC or ACC.** Panels on the left show CCMs (color code depicts the Pearson correlation coefficients between vectors of t-statistics associated with regression coefficient estimates for the $k^{th}$ attended cue at cue onset time $t$, across integration units, going from blue – low correlation – to pink – high correlation –), organized by cue index in the sequence ($k$=1, 2 or 3) and response time ($t$=1, 2 or 3). Asterisks indicate significant correlations, with p-value < 0.0007 (correction for multiple comparisons across

CCM cells). Panels on the right show CCM cells from monkey M (y-axis) plotted against CCM cells from monkey F (x-axis).

Second, Figure S3 shows the raw CCMs of each monkey and pooled recorded neurons of both monkeys together, in either the OFC, dlPFC or ACC.

Here again, OFC, dlPFC and ACC neurons seem to exhibit regionally specific representational geometries.

Strikingly, both RDM and CCM metrics also show inter-individual differences. This raises the question of whether inter-regional differences are greater than inter-individual differences, or not. To test this, we compute the average log-ratio of RDM and CCM variance across regions and across subjects, where the average is taken over the entries of RDM and CCM matrices. We obtain the distribution of the log-ratio under the null using 10,000 permutations of region labels. Figure S4 below summarizes this analysis.

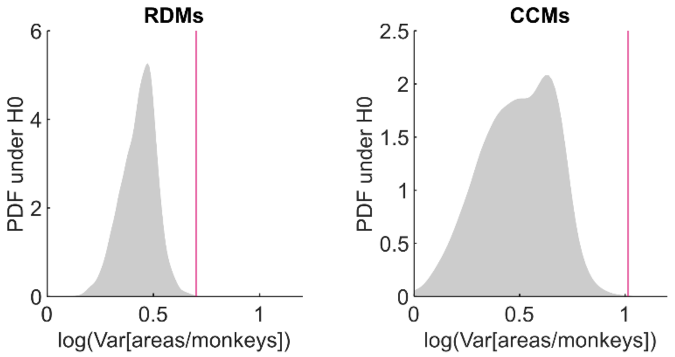


**Fig. S4 | Comparison of RDM and CCM variance over brain regions and over subjects.** Grey areas depict the distribution, under the null, of the log ratio of RDM (left) and CCM (right) variance over regions and monkeys. Pink lines show the log ratio of variances in the actual data.

The ensuing p-values are below $5 \times 10^{-4}$, i.e., inter-regional differences are significantly stronger than inter-individual differences.

#### 3. Do irrational RNNs accurately predict inter-individual differences in OFC's CCMs?

In the main text, we show that irrational RNN training does improve the resemblance to patterns of activity within the OFC, when compared to rational RNNs. But do irrational RNNs accurately predict inter-individual differences in the informational geometry of the OFC?

To test this, we asked whether CCM cells of RNNs tend to align with CCM cells in the OFC, across monkeys. That is, we quantify the average regression slope between predicted and actual CCM entries or cells (across monkeys). We obtain the distribution of the average slope (over CCM cells) under the null using 10,000 permutations of monkeys' labels. The results of this analysis are summarized in Figure S5 below.

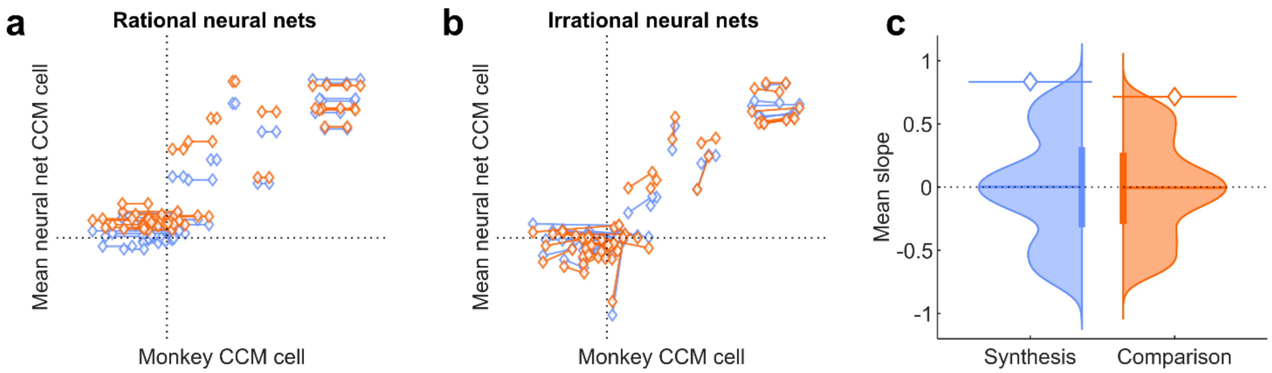


**Fig. S5 | Comparison of CCM entries across monkeys. a**, The CCM entries of rational RNNs (y-axis) are plotted against the CCM entries of their associated monkey (x-axis). Each dot is a given CCM entry (blue: value synthesis, orange: value comparison), and lines relate pairs of entries across monkeys. Accurate predictions of inter-individual differences would show up as oblique lines, aligned with the main diagonal (positive slopes). **b**, Same thing for re-trained (irrational) RNNs. **c**, Distribution of the average slope (across CCM cells) under the null, for the irrational value synthesis RNNs (blue) and irrational value comparison RNNs (orange). The horizontal lines with a diamond show the average slope in the actual data.

Panel b in Fig. S5 shows that retrained RNNs make subject-specific CCM predictions that tend to align with their corresponding CCM entries in the OFC. As expected, this is not the case for rational RNNs, since their predicted CCM entries are (by construction) invariant across monkeys. This analysis demonstrates that predicted inter-individual differences in CCM cells are significantly accurate for both types of retrained RNNs (synthesis models: $p = 3\times10^{-3}$; comparison models: $p = 1\times10^{-2}$).

### 4. Are decisions triggered later in time more difficult?

In the main text, we show that retrained (irrational) RNNs accumulate perturbations in their value computations over decision time and we compare this prediction to the rate of monkeys' irrational choices as a function of the time at which choices are triggered. Although we find that monkeys also tend to become more irrational for choices that are triggered later in time, this effect may be partially driven by the fact that more difficult choices may require more decision-relevant information and may thus be triggered later in time. The issue here is that more difficult choices would also tend to be more irrational on average, since small variations in valuations have a greater chance to yield preference reversals. To check this, we evaluated the average choice ease, measured in terms of the absolute subjective value difference between options, at each choice onset time. Figure S6 below shows choice ease as a function of choice onset time, for both *option* and *attribute* trials and each monkey.

Although decisions that are triggered later in time are more difficult for *attribute* trials (Monkey F, two-sample two-sided t-test between step 2 and step 4: $t(2755) = 7.0$, $p = 3\times10^{-12}$, Cohen's $d = -0.35$; Monkey M: $t(3639) = 12$, $p < 10^{-15}$, Cohen's $d = 1.00$) this is the reverse for *option* trials (Monkey F: $t(2730) = 22$, $p < 10^{-15}$, Cohen's $d = 1.00$; Monkey M: $t(4316) = 21$, $p < 10^{-15}$, Cohen's $d = 1.58$). Nevertheless, there is a clear variation of choice difficulty across decision times, which is why we correct for this effect when analyzing the rate of irrational choices across choice onset times

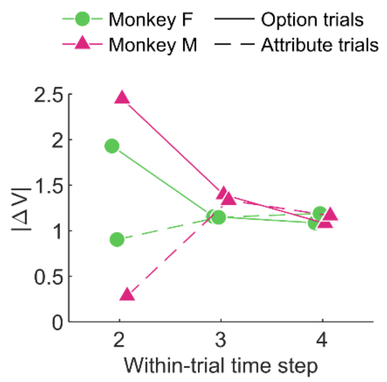


**Fig. S6 | Choice ease as a function of choice onset time.** Average choice ease, as measured in terms of the absolute value difference between options (where option values are derived from each monkey's estimated value profile), is plotted against choice onset time, for both *option* (solid line) and *attribute* (dashed line) trials. Colors and marker shapes designate monkeys (green circle: monkey F, pink triangle: monkey M).

### 5. How robust is our model validation approach?

In the main text, we report many results on irrational RNNs, where irrational RNNs have been obtained by retraining (fine-tuning) the recurrent connections of rational RNNs to explain monkeys' irrational choices. In particular, we show that such irrational RNNs make neural predictions that are closer to OFC neurons than their rational counterparts. Importantly, we demonstrate that these predictions are specific to the OFC, in the sense that they are less resemblant to either dlPFC or ACC neural activity patterns. But do these results hold if we derive irrational RNNs by training them directly on monkeys' choices, without initializing them using trained rational RNNs?

To check this, we repeated the RNN training procedure (1,000 instances in each RNN cohort), this time directly relying on monkeys' choices at decision onset times. Note that this "direct" procedure produces RNNs with poorer across-trials generalization, compared to the fine-tuning procedure used in the main text. We then compared the activity patterns of these irrational RNNs to activity patterns in the OFC, the dlPFC and the ACC exactly as we did before. The results of these analyses are summarized on Figure S7 below.

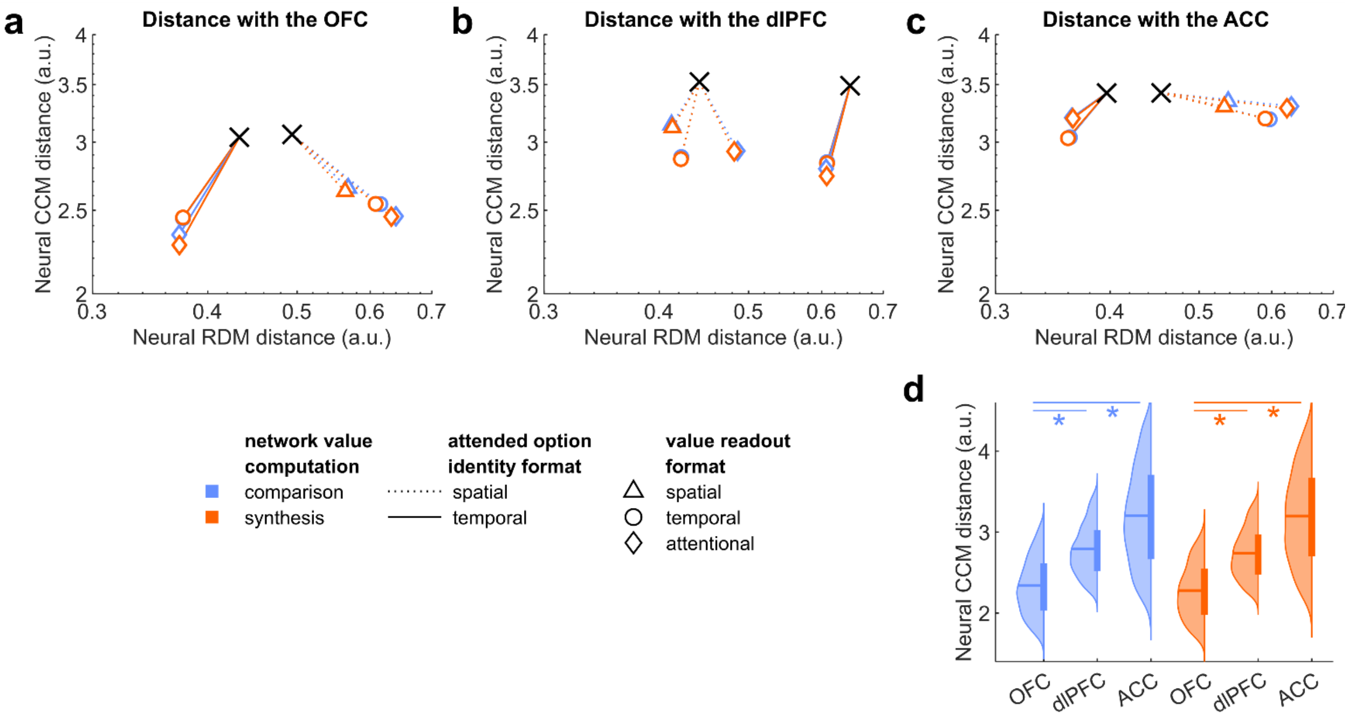


**Fig. S7 | Evaluating the accuracy of neural predictions of irrational RNNs directly trained on monkeys' choices.** **a**, Neural distance trajectories between OFC and RNN cohorts during direct irrational training. Points show the average distance between activity patterns of OFC recordings and RNN models (across the 1,000 RNN instances), computed using either RDMs (x-axis) or CCMs (y-axis) metrics, either at initialization (black crosses) or at training convergence (see legend). **b** and **c**, Same, for dlPFC and ACC neurons, respectively. **d**, Distributions (across cohort instances) of final neural CCM distances between irrational models and the OFC, the dlPFC and the ACC, for value synthesis (blue) and value comparison (orange) RNNs that rely on a temporal option identity format and an attentional value readout format. Asterisks indicate significant differences between distances with OFC on the one hand, and both dlPFC and ACC on the other hand, with p-value < 0.005. This panel reproduces Fig 3g in the main text.

Strikingly, irrational RNNs that are directly trained on monkeys' choices make neural predictions that are similarly specific to the OFC. Importantly, the two irrational cohorts that are most resemblant to the OFC are also the same as selected rational models in the main text, i.e. RNNs that operate either value synthesis or value comparison under a temporal option identity format and an attentional value readout format. This means that the results of our model validation approach do not depend upon how we derive irrational RNNs.

### 6. How specific is the increased tolerance of irrational RNNs?

In the main text, we report that irrational RNNs are more tolerant to neural loss than their rational counterpart. We obtained this result by silencing units in the integration layer, and quantifying the retained rate of rational choices while increasing the lesion extent. Here, we ask whether irrational RNNs are also more tolerant to 1) internal disconnections and 2) neural noise.

To test the tolerance to internal disconnections, we consider a proportion $p \in \{0.1, 0.2, \dots, 1\}$ of lesioned connections. Let $x_1 = 9$ and $x_2 = 10$ denote the number of units in the first and second hidden layers, respectively. Each RNN therefore comprises $x_1 \times x_2 = 90$ feedforward connections and $x_2 \times x_2 = 100$ recurrent connections. Lesions are distributed proportionally across the two connection sets $\boldsymbol{W}_{encode}$ and $\boldsymbol{W}_{recurrent}$, such that $n_1 \in \{9, 18, \dots, 90\}$ feedforward connections and $n_2 \in \{10, 20, \dots, 100\}$ recurrent connections are lesioned. A specific lesion combination is represented by lesion maps $\boldsymbol{C}_p^1 \in \mathbb{R}^{9\times10}$ and $\boldsymbol{C}_p^2 \in \mathbb{R}^{10\times10}$. Artificial lesions are implemented by setting the corresponding lesion weights to 0. Let $z_{model}\left(\boldsymbol{U}, t, \boldsymbol{C}_p^1, \boldsymbol{C}_p^2\right) \in \{0, 1\}$ denote the RNN's simulated choice at time $t$ in response to an input sequence $\boldsymbol{U}$, under lesion maps $\boldsymbol{C}_p^1$ and $\boldsymbol{C}_p^2$. Let $z_{rational}(\boldsymbol{U}, t)$ denote the rational choice (i.e., the preferred option based upon options' expected value) given the same input sequence. We define the tolerance to connection lesions $\overline{T_C}$ as the retained rate of rational choice, averaged over 100 randomly sampled pairs of lesion maps for each lesion level between 10% and 50%:

$$\overline{T_C} = \frac{1}{5 \times 100 \times |U| \times T} \sum_{p=0.1}^{0.5} \sum_{i=1}^{100} \sum_{\boldsymbol{U}} \sum_{t=1}^{T} 1_{\{z_{model}(\boldsymbol{U},t,\boldsymbol{C}_p^1,\boldsymbol{C}_p^2)=z_{rational}(\boldsymbol{U},t)\}}$$

where $i$ indexes independently sampled lesion map pairs.

To test the tolerance to neural noise, we added noise to integration units' activations. Recall the standard propagation equation:

$$\boldsymbol{X}_2(t) = f\big(\boldsymbol{W}_{forward}.\boldsymbol{X}_1(t) + \boldsymbol{W}_{recurrent}.\boldsymbol{X}_2(t-1) - \boldsymbol{B}_2\big)$$

where $\boldsymbol{X}_1(t) \in \mathbb{R}^9$ and $\boldsymbol{X}_2(t) \in \mathbb{R}^{10}$ denote the unit activation vectors of the first and second hidden layers at time $t$, respectively, and $\boldsymbol{B}_2 \in \mathbb{R}^{10}$ is the bias vector applied to the second hidden layer (see Methods).

To model neural noise, the equation is modified as follows:

$$\boldsymbol{X}_2(t) = f\big(\boldsymbol{W}_{forward}.\boldsymbol{X}_1(t) + \boldsymbol{W}_{recurrent}.\boldsymbol{X}_2(t-1) - \boldsymbol{B}_2\big) + \boldsymbol{\varepsilon_\sigma}$$

where $\boldsymbol{\varepsilon_\sigma} \sim \mathcal{N}(0, \sigma I)$ is a noise vector whose elements are independently drawn from a centered normal distribution with variance $\sigma$. Let $z_{model}(\boldsymbol{U}, t, \sigma) \in \{0, 1\}$ denote the RNN's simulated choice at time $t$ in response to an input sequence $\boldsymbol{U}$, under neural noise of variance $\sigma$. We define the tolerance to neural noise $\overline{T_N}$ as the retained rate of rational choice, averaged over 50 independent noise realization for each variance level $\sigma \in \{0.001, 0.005, 0.01, 0.05, 0.1, 0.5\}$:

$$\overline{T_N} = \frac{1}{6 \times 50 \times |U| \times T} \sum_{\sigma} \sum_{i=1}^{50} \sum_{\boldsymbol{U}} \sum_{t=1}^{T} 1_{\{z_{model}(\boldsymbol{U},t,\sigma) = z_{rational}(\boldsymbol{U},t)\}}$$

The results of all three tolerance analyses are detailed in Figure S8 below.

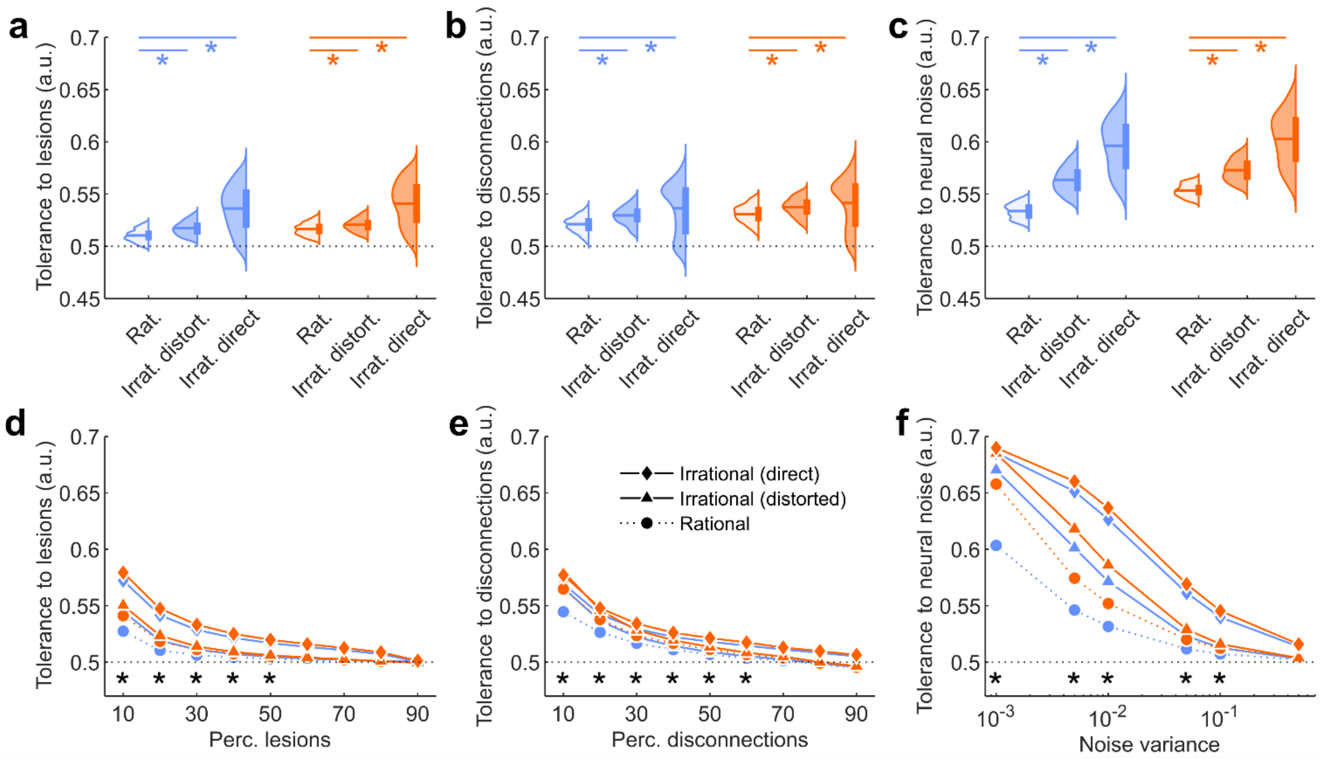


**Fig. S8 | Comparing the tolerance to structural lesions and neural noise in rational and irrational RNNs. a**, Distributions (across network instances) of tolerance to unit lesions, measured as the retained rate of rational choice from 10% to 50% of lesions, for value synthesis (blue) and value comparison (orange) RNNs that rely on a temporal option identity format and an attentional value readout format. Asterisks indicate a significant difference between rational (light) and irrational (dark) RNNs, with p-value < 0.005. Note that both types of irrational models (either retrained from rational RNNs or directly trained from monkeys' irrational choices) are shown here. **b** and **c**, Same format as panel **a**, showing the tolerance to connection lesions from 10% to 50% of lesions (**b**), and the tolerance to neural noise averaged the over the full range of noise variance (**c**). **d**, Tolerance to unit lesions as a function of lesion level. Asterisks indicate significant differences between rational (dotted line, circles) models and irrational (solid lines) models, including both retrained (triangles) or trained directly (diamonds) RNNs, for value synthesis (blue) and value comparison (orange) models, with p-value < 0.005 for each comparison. **e** and **f**, Same format as panel **d**, showing the tolerance to connection lesions (**e**) and neural noise (**f**).

In brief, irrational RNNs exhibit significantly stronger tolerance to both disconnections and neural noise than their rational counterparts, irrespective of value computations. This holds true both for irrational RNNs that are retrained from rational RNNs (disconnections, synthesis models, two-sample two-sided t-test: $t(1999) = 23$, $p < 10^{-15}$, Cohen's $d = 0.52$; comparison models: $t(1999) = 19$, $p < 10^{-15}$, Cohen's $d = 0.43$; neural noise, synthesis models: $t(1999) = 76$, $p < 10^{-15}$, Cohen's $d = 1.69$; comparison models: $t(1999) = 59$, $p < 10^{-15}$, Cohen's $d = 1.29$) and for irrational RNNs that are directly trained from monkeys' choices (disconnections, synthesis models, two-sample two-sided t-test: $t(1999) = 21$, $p < 10^{-15}$, Cohen's $d = 0.48$; comparison models: $t(1999) = 15$, $p < 10^{-15}$, Cohen's $d = 0.34$; neural noise, synthesis models: $t(1999) = 86$, $p < 10^{-15}$, Cohen's $d = 1.93$; comparison models: $t(1999) = 70$, $p < 10^{-15}$, Cohen's $d = 1.57$).
